# A novel adenovirus 19K/IX protein enhances infection by attenuating proteasomal degradation of tyrosine-ubiquitinated capsid protein IX

**DOI:** 10.64898/2026.04.24.720336

**Authors:** Erik Schubert, Vasileios Rafail Evangelopoulos, Mårten Larsson, Tanel Punga

## Abstract

Human adenoviruses (HAdVs) are double-stranded DNA viruses that cause a wide range of diseases, including respiratory, ocular, and gastrointestinal infections. HAdVs rely extensively on alternative splicing to expand their coding capacity and regulate gene expression during infection. Using replication-competent epitope-tagged HAdV-C5 viruses, we investigated the function of a newly identified alternatively spliced viral transcript, 19K/IX, derived from the E1B19K and pIX genes. We show that the 19K/IX protein binds to and stabilizes the minor capsid protein IX (pIX), thereby attenuating its proteasomal degradation and enhancing the production of infectious viral progeny. Mechanistically, we demonstrate that pIX degradation is mediated by non-canonical ubiquitination of the conserved tyrosine residues Y14 and Y49, which confer pIX subnuclear localization and partially mediate its interaction with the PSMC3 subunit of the 26S proteasome. Collectively, our study identifies 19K/IX as a novel regulator of HAdV-C5 infection and suggests that non-canonical tyrosine ubiquitination contributes to the regulation of capsid protein IX stability.

**Importance:** Human adenoviruses (HAdVs) are clinically important pathogens and widely used as therapeutic tools. Because of their compact genomes, HAdVs use their genetic information very efficiently through alternative pre-mRNA processing. Here, we describe and characterize 19K/IX, a novel fusion protein generated by alternative splicing of two viral genes during infection. We demonstrate that 19K/IX stabilizes capsid protein IX by preventing its degradation, revealing a previously unrecognized strategy by which HAdVs ensure efficient production of new virus particles. These findings deepen our understanding of HAdV-C5 gene regulation and protein homeostasis and highlight the functional importance of non-canonical HAdV transcripts.

## Introduction

Human adenoviruses (HAdVs) are globally prevalent pathogens that cause respiratory, ocular, and gastrointestinal diseases (1, 2). While most HAdV infections resolve without complications, certain highly virulent HAdV types, such as HAdV-B3, HAdV-E4, HAdV-B7, and HAdV-B55, can have severe or fatal outcomes (1, 3–5). In addition, HAdV-C2 and HAdV-C5, considered low-pathogenic in immunocompetent individuals, occasionally cause life-threatening diseases in immunocompromised individuals (6, 7).

To accommodate its limited genome size, HAdV pre-mRNAs undergo extensive alternative splicing and polyadenylation, generating multiple viral transcripts that encode a wide range of viral regulatory and structural proteins (8–11). Advanced long-read RNA sequencing has revealed the extreme complexity of the transcriptomes of the genetically closely related HAdV-C2 and HAdV-C5 and identified multiple novel, potentially protein-encoding viral transcripts (12–15). One newly identified transcript, 19K/IX, is generated by alternative splicing of the viral early E1B pre-mRNA (schematically shown in Fig. 1A) and is derived from the E1B transcription unit (12–14). The latter is driven by its own promoter and generates multiple transcripts, including transcripts encoding the anti-apoptotic E1B19K protein and the p53 antagonist E1B55K, which are encoded by overlapping open reading frames (16). Curiously, the distal region of the E1B transcription unit also contains the gene encoding protein IX (hereafter as pIX), which uses the same cleavage and polyadenylation signal (CPAS) to define the 3′ ends of both the E1B and pIX transcripts (14). Notably, the 5′ portion of the E1B19K coding sequence can be alternatively spliced to the pIX coding sequence, generating an in-frame 19K/IX fusion transcript. The pIX gene is expressed independently from its own promoter, generating pIX (17, 18), a multifunctional minor capsid “cement” protein that contributes to capsid stabilization by binding to the viral capsid protein hexon (19–21). It has also been shown that pIX activates the HAdV early (E1A) and late (major late transcription unit, MLTU) promoters in transiently transfected cells (22–24). In addition, pIX is implicated in host nuclear reorganization, as its expression leads to the formation of specific nuclear inclusions (22, 24–26). Although pIX is dispensable for efficient virus growth, its absence reduces infectivity and slows virus growth (24, 27, 28).

**Fig 1.**
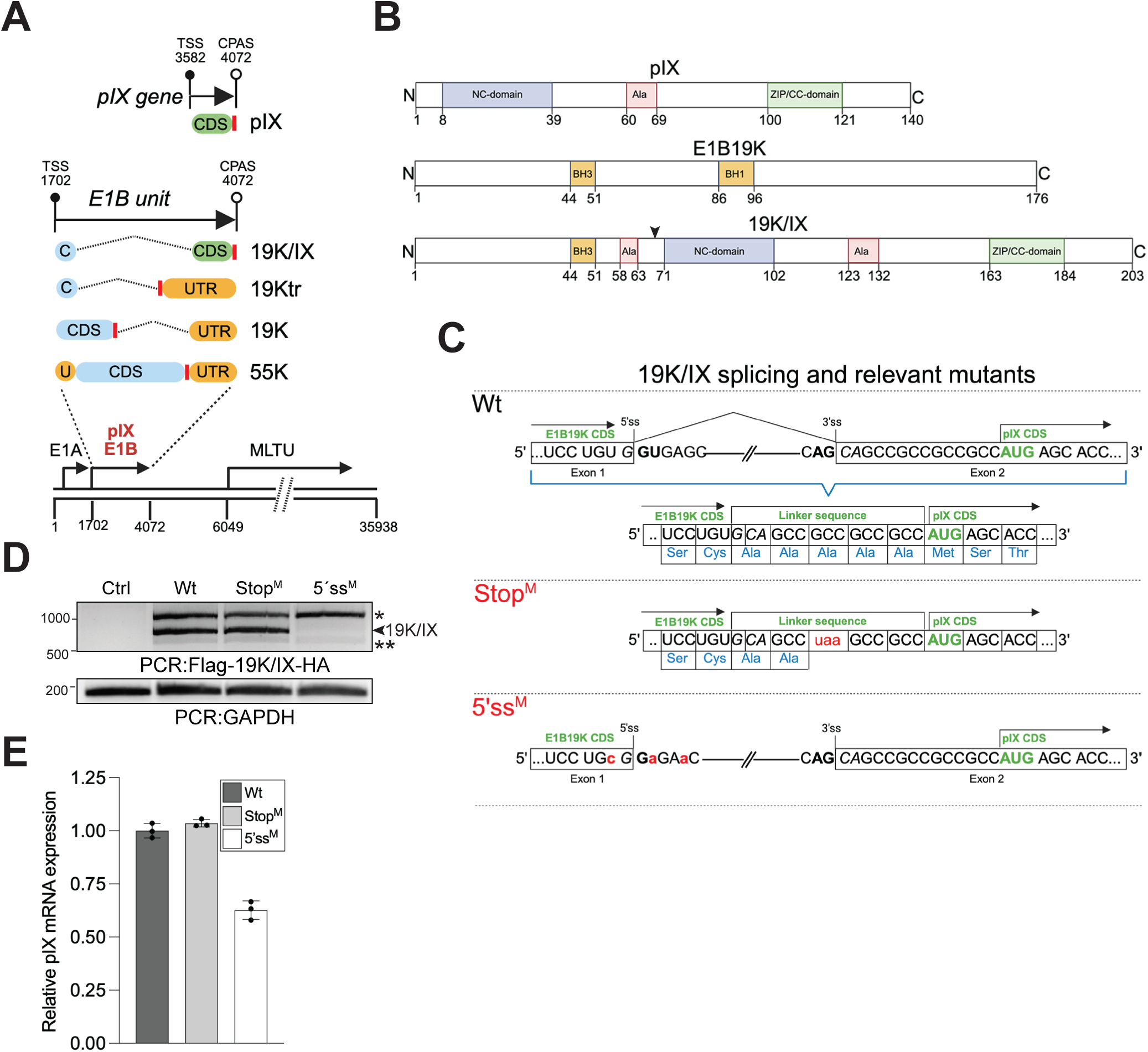
Origin of 19K/IX transcript and its mutant viruses. **(A)** Simplified schematic of the E1B transcription unit and the pIX gene in the HAdV-C5 genome (AC_000008.1), adapted from (14). Light blue blobs; E1B coding DNA sequence (labeled as C or CDS), green blob; pIX CDS, yellowish blobs; untranslated regions (labeled as U or UTR), the vertical red line; translation termination signal (TTS), horizontal arrow; indicated gene unit, filled lollipop; transcription start site (TSS), opened lollipop; the cleavage and polyadenylation signal (CPAS), MLTU; major late transcription unit, 55K; E1B55K, 19K; E1B19K, 19Ktr; truncated isoform of E1B19K. The naming of E1B transcripts and virus genome coordinates is based on (14). **(B)** A schematic overview of the pIX, E1B19K, and 19K/IX proteins. NC-domain; N-terminal conserved domain, Ala; alanine-rich sequence, ZIP/CC-domain; putative leucine zipper/coiled-coil domain (23, 24), BH3 and BH1; Bcl-2 homology 1 and 3 (52). A vertical arrowhead indicates the fusion point between E1B19K and pIX. **(C)** A schematic overview of the splicing pattern between the E1B19K and pIX CDS to generate the 19K/IX transcript. Wt; splicing between E1B19K cryptic 5’ splice site (5’ss, nucleotide (nt) 175 of the 528 nt E1B19K CDS) and pIX 3’ splice site (3’ss) (upper panel), Stop^M^; splicing pattern as in Wt, but with an in-frame translation termination codon (uaa) intoduced between the E1B19K and pIX CDS, terminating 19K/IX translation (middle panel), 5’ss^M^; no splicing between E1B19K 5’ss and the pIX 3’ss due to point mutations in the E1B19K 5’ss (lower panel). Linker sequence; nucleotide sequence encoding five alanine residues generated by splicing of the E1B19K cryptic 5’ss to the 3’ss upstream of the pIX CDS, red; introduced mutations in Stop^M^ and 5’ss^M^, light blue; amino acids. **(D)** Endpoint PCR amplification of the Flag-19K/IX-HA transcript in infected A549 cells (24 hpi). The asterisks indicate aberrant transcripts that will not encode the 19K/IX protein. PCR was performed using primers annealing to the 5′ Flag-tag and 3′ HA-tag sequences (PCR: Flag-19K/IX-HA), which were integrated into the HAdV-C5 genome (Fig. 2A), GAPDH was used as the amplification control. **(E)** Relative pIX mRNA expression detected with qRT-PCR from samples in panel D. Primers anneal to the central region of the pIX CDS, pIX expression was normalized to HRPT1. Data are from one independent experiment (n = 1) performed in technical triplicate.

The 19K/IX transcript is abundant in infected cells (13, 14). However, the expression of the 19K/IX protein during infection and its potential role in the viral life cycle remain unknown. Here, we address these questions by characterizing the expression and function of 19K/IX during HAdV-C5 infection.

## Results

### Origin of 19K/IX transcript and its mutant viruses

To determine the role of 19K/IX during infection, we generated two replication-competent HAdV-C5 viruses with mutations that disrupt 19K/IX expression. The first mutant virus (referred to as Stop^M^) was generated by converting an alanine codon in the linker sequence between the E1B19K and pIX coding DNA sequence (CDS) into a stop codon (UAA), thereby selectively disrupting translation of the 19K/IX mRNA. In the second mutant virus (referred to as 5’ss^M^), a cryptic 5’ splice site (ss) within the E1B19K CDS was mutated, assuming that this mutation would not allow splicing to the alternative 3’ ss present upstream of the pIX CDS and thereby preventing 19K/IX mRNA production (Figs. 1B and 1C). As expected, the 19K/IX transcript was not detected in 5’ss^M^-infected human lung epithelial cells (A549), but was present in Stop^M^-infected cells (Figs. 1D and 1E). To visualize the 19K/IX protein, we modified HAdV-C5 genomes by inserting a Flag antibody epitope tag at the 5’ end of the E1B19K CDS and an HA antibody epitope tag at the 3’ end of the pIX CDS (Fig. 2A). This tagging strategy enables the detection of E1B19K (Flag staining), pIX (HA staining), and the 19K/IX protein (Flag and/or HA staining) in cells infected with antibody epitope-modified HAdV-C5 (referred to as Wt in this study).

**Fig 2.**
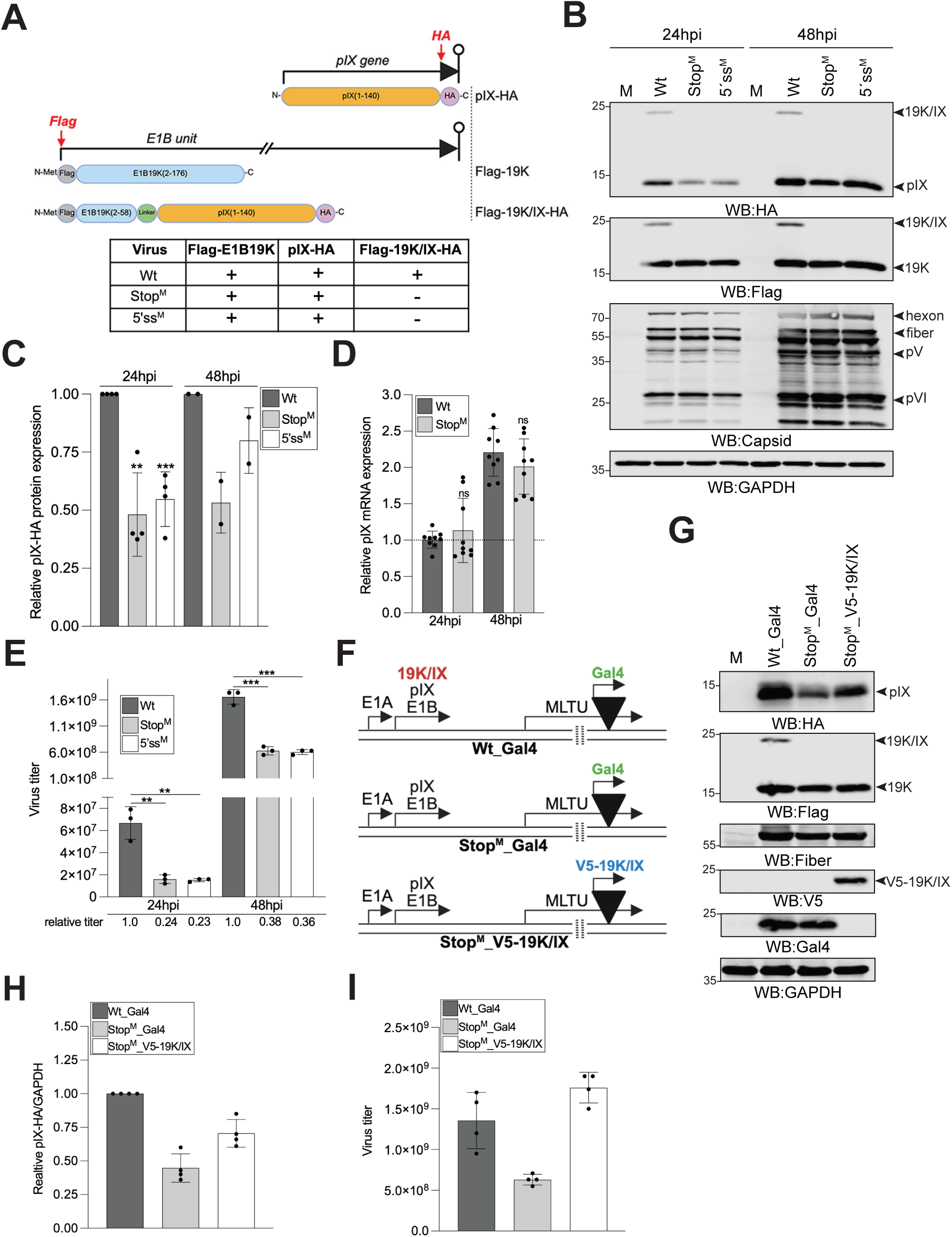
Lack of the 19K/IX protein reduces pIX accumulation and viral progeny production. **(A)** Graphical overview of the pIX-HA, Flag-E1B19K, and Flag-19K/IX-HA proteins. Integration of the Flag antibody epitope tag sequence at the 5’ end of the E1B transcription unit and the HA antibody epitope tag sequence at the 3’ end of the pIX gene is shown with vertical red arrows. Protein lengths in amino acids are indicated in parentheses. The inset table summarizes which tagged proteins are expressed in epitope-tagged viruses. **(B)** A549 cells were infected with Wt, Stop^M^, or 5’ss^M^ viruses. Whole cell lysate (WCL) was prepared at 24 hpi and 48 hpi and analyzed by western blotting (WB) with the anti-HA, anti-Flag (also detects Flag-E1B19K (referred to as 19K)), anti-capsid, and anti-GAPDH antibodies. A representative WB is shown. **(C)** Relative quantification of independent experiments (n ≥ 2) from panel B. **(D)** Relative pIX mRNA expression after normalization to HRPT1. Quantification of three independent experiments (n = 3) analyzed in technical triplicates. **(E)** Virus titer (FFU/ml) was determined by re-infecting A549 cells with Wt, Stop^M^, and 5’ss^M^ virus-containing cell lysates. The relative virus titer is shown after normalizing the Wt samples at 24 hpi and 48 hpi to 1, (n = 3). **(F)** A schematic overview of the Wt_Gal4, Stop^M^_Gal4, and Stop^M^_V5-19K/IX viruses. A vertical arrowhead indicates the position at the E3 region where the CMV-Gal4-poly(A) or CMV-V5-19K/IX-poly(A) expression cassettes were inserted. A horizontal arrow indicates the relative location of the MLTU on the viral genome. **(G)** A549 cells were infected with the Wt_Gal4, Stop^M^_Gal4, or Stop^M^_V5-19K/IX at MOI=5. WCL was prepared at 48 hpi and analyzed by WB with the indicated antibodies. A representative WB is shown. **(H)** Relative accumulation of pIX-HA protein, normalized to GAPDH, from two independent experiments (n = 2) in technical duplicates, from panel G. **(I)** Virus titer (FFU/ml) determined by re-infecting A549 cells with Wt_Gal4, Stop^M^_Gal4, or Stop^M^_V5-19K/IX virus-containing cell lysates. Virus titer was determined as in panel E from two independent experiments (n = 2) in technical duplicate.

### Lack of the 19K/IX protein reduces pIX accumulation and production of viral progeny

We first validated 19K/IX expression during infection using our epitope-tagged viruses (Fig. 2A). Consistent with transcriptomics data (14), the 19K/IX protein was readily detected from 16 to 48 hours post-infection (hpi) in A549 cells using Flag and HA antibodies (Fig. 2B and Fig. S1A). In contrast, the 19K/IX protein was not detected in the 5’ss^M^- and Stop^M^-infected cells, confirming that both mutations (Fig. 1C) indeed eliminated 19K/IX expression (Fig. 2B). The absence of the 19K/IX protein did not detectably affect the expression of the early E1B19K or late viral capsid proteins (Fig. 2B). Notably, we observed reduced accumulation of the pIX protein in 5’ss^M^- and Stop^M^-infected cells, particularly at 24 hpi (Figs. 2B and 2C, Fig. S1A). The pIX mRNA levels were not significantly reduced in Stop^M^-infected cells (Fig. 2D), suggesting that lower pIX levels (Fig. 2C) were mainly due to reduced protein synthesis or degradation. Given the established role of pIX in virus capsid stabilization (19–21), we examined whether reduced pIX levels affected viral progeny production. Infectious viruses isolated from the Stop^M^- and 5’ss^M^-infected cells revealed reduced virus titers, implying that 19K/IX enhances production of infectious progeny (Fig. 2E). To confirm further the 19K/IX effect on pIX expression, we generated a replication-competent virus with a CMV promoter-driven V5-epitope-tagged 19K/IX CDS inserted into the E3 region of the Stop^M^ virus (referred to as Stop^M^_V5-19K/IX, Fig. 2F). As a control, a CMV-driven yeast transcription factor Gal4 DNA-binding domain (Gal4) was inserted into the same E3 region of the Wt and Stop^M^ (referred to as Wt_Gal4 and Stop^M^_Gal4, respectively). Expression of V5- 19K/IX enhanced pIX accumulation in Stop^M^_V5-19K/IX-infected cells, whereas Gal4 expression did not have the same effect in Stop^M^_Gal4-infected cells (Figs. 2G and 2H). Furthermore, expression of V5-19K/IX restored Stop^M^_V5-19K/IX virus titers to similar levels to those observed for the Wt_Gal4 virus (Fig. 2I). Together, these data demonstrate that 19K/IX expression enhances pIX accumulation and production of infectious virus progeny.

### The 19K/IX protein interacts with pIX to form an oligomeric complex

The finding that 19K/IX regulates pIX accumulation (Fig. 2B) led us to investigate whether the two proteins interact. To this end, we performed an anti-Flag immunoprecipitation (IP) to isolate the Flag-19K/IX-HA protein from Wt-, Stop^M^-, and 5’ss^M^-infected A549 cells. Because all three viruses express the Flag-E1B19K protein (Fig. 2B), this protein was also recovered in the anti-Flag IP. Notably, a strong interaction with pIX-HA was detected in Wt-infected cells expressing Flag-19K/IX (Fig. 3A). The Flag IP experiments were repeated in transiently transfected A549 cells with the plasmids expressing the V5-tagged 19K/IX (V5-19K/IX), Flag-tagged pIX (Flag-pIX), and HA-tagged pIX (HA-pIX). Similar to virus infection, the Flag-pIX interacted with the V5-19K/IX protein (Fig. 3B, lanes 8 and 9, marked with an arrowhead). Previous studies have shown that pIX self-associates into dimeric/oligomeric forms (24, 29). We therefore asked whether transient V5-19K/IX expression influences pIX-pIX formation. To test this, we co-expressed pIX with different epitope tags (Flag-pIX and HA-pIX) and assessed their mutual interaction as the oligomerization readout. Although Flag-pIX and HA-pIX co-immunoprecipitated (Fig. 3B, lane 7), confirming complex formation, co-expression of V5-19K/IX did not detectably affect pIX-pIX formation under our experimental conditions (Fig. 3B, lane 8). Interestingly, transient V5-19K/IX expression increased HA-pIX and Flag-pIX protein accumulation, suggesting that 19K/IX may enhance pIX stability in A549 cells (Fig. 3B, lanes 1 to 4). To confirm the pIX and 19K/IX oligomerization during infection, we treated A549 cells with the membrane-permeable, non-cleavable protein crosslinker disuccinimidyl suberate (DSS) to stabilize and detect distinct oligomeric pIX and 19K/IX species. We detected pIX as a homodimer (pIX-pIX) as well as a heterodimer with the 19K/IX protein (pIX-19K/IX) in Wt-infected cells in a DSS concentration-dependent manner (Fig. 3C). In Stop^M^-infected cells, the pIX-19K/IX complex was not detected, and pIX-pIX formation was reduced (Fig. 3C), consistent with lower pIX expression in these cells (Fig. 2B). To assess their spatial relationship, we analyzed 19K/IX and pIX in nuclear, cytosolic, and mitochondria-enriched fractions in A549 cells (30). Notably, pIX was detected predominantly in the cytosolic fraction, with no drastic differences between Wt-, Stop^M^-, and 5′ss^M^-infected cells (Fig. 3D, lanes 1, 4, and 7). The 19K/IX protein was likewise enriched in the cytosol and was detected in all three fractions (Fig. 3D, lanes 1-3). In contrast, full-length E1B19K and its recently identified truncated isoform, 19Ktr (14), were almost exclusively detected in the mitochondria-enriched fraction (Fig. 3D, lanes 2, 5, and 8). Previous studies have shown that pIX expression induces distinct nuclear inclusions in both transfected and virus-infected cells (22, 24–26), which is consistent with our observations (Figs. S1B, S1C and S1D). We therefore asked whether 19K/IX expression alters the pattern of pIX-specific nuclear inclusions. As shown in Fig. 3E, V5-19K/IX colocalized with pIX-HA in nuclear inclusions, with colocalization also in the cytoplasm, without visually detectable alteration of the pIX nuclear inclusions in virus-infected cells. Notably, transient expression of 19K/IX alone did not induce nuclear inclusions (Figs. S1C, S1D and S1E). This observation suggests that pIX may contribute to the relocalization of 19K/IX to the nuclear inclusions.

**Fig 3.**
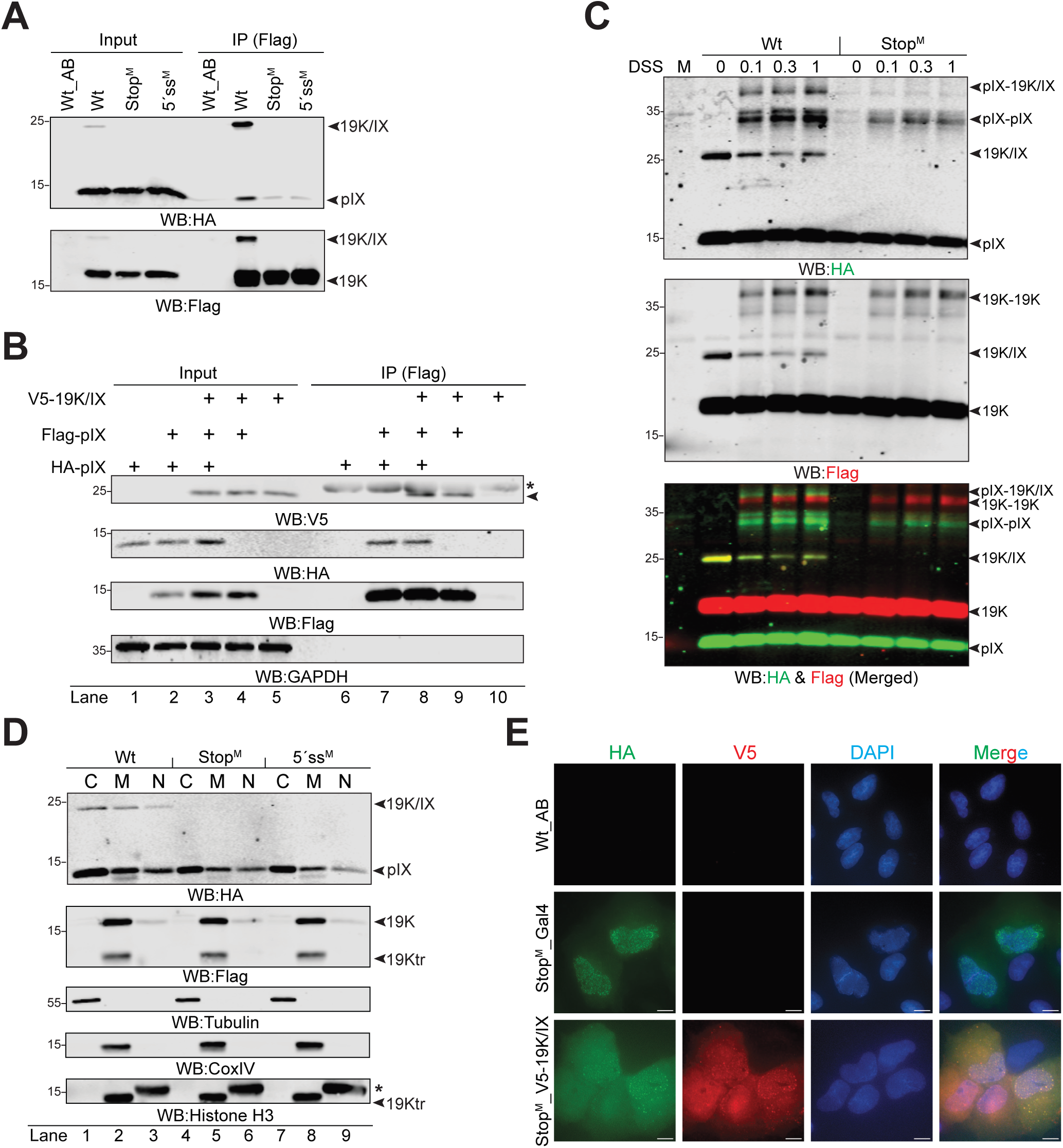
The 19K/IX protein interacts with pIX to form an oligomeric complex. **(A)** 19K/IX interacts with pIX in Wt-, Stop^M^- or 5’ss^M^-infected A549 cells. Note that anti-Flag immunoprecipitation (IP) recovers both Flag-19K/IX-HA and Flag-E1B19K. Wt_AB; unmodified AdenoBuilder-based wild-type HAdV-C5 expressing no epitope-tagged proteins. **(B)** V5-tagged 19K/IX interacts with the oligomeric pIX. Anti-Flag IP from A549 cells transfected with plasmids encoding V5-19K/IX, Flag-pIX or HA-pIX. Proteins were detected by WB using the indicated antibodies. An asterisk; anti-Flag IgG light chain, an arrowhead; V5-19K/IX. Lane numbers are shown below the images. **(C)** 19K/IX oligomerizes with pIX. Wt- or Stop^M^-infected A549 cells at 24 hpi were treated with DSS crosslinker at different concentrations (0.1-1mM). Predicted protein monomers, homodimers, and heterodimers are indicated. M; mock, non-infected cells. Flag-E1B19K (19K) monomers and dimers are also indicated due to Flag antibody staining. **(D)** Biochemical fractionation of the A549 cells infected with Wt, Stop^M,^ or 5’ss^M^ at 24 hpi. Cytosolic (C), mitochondria-containing (M), and nuclear (N) fractions were analyzed by WB using antibodies against HA, Flag, Tubulin (cytosolic marker), CoxIV (mitochondrial marker), and histone H3 (nuclear marker). An asterisk indicates migration of histone H3. Residual Flag antibody-stained 19Ktr protein is visible on the histone H3-stained membrane (marked with an arrowhead). Lane numbers are shown below the images. **(E)** 19K/IX and pIX colocalize in nuclear inclusions. Indirect immunofluorescence (IF) in A549 cells infected with Wt_AB (unmodified AdenoBuilder-based wild-type HAdV-C5 expressing no epitope-tagged proteins), Stop^M^_Gal4, or Stop^M^_V5-19K/IX viruses at 24 hpi. Proteins were stained with the HA and V5 antibodies, and nuclei were stained with DAPI. Scale bar = 10 µm.

### 19K/IX counteracts pIX degradation in a proteasome-dependent, lysine-independent manner

The observation that transient 19K/IX expression increased pIX levels (Fig. 3B) suggested that 19K/IX may regulate pIX stability. To test this hypothesis, we performed a cycloheximide (CHX) chase experiment (31), which blocks *de novo* protein synthesis, in Wt- or Stop^M^-infected A549 cells and monitored pIX accumulation over 24 h. Notably, pIX was significantly more resistant to degradation in Wt-infected cells compared to Stop^M^-infected cells (Figs. 4A and 4B). To test whether pIX turnover is proteasome-dependent, cells were treated with the proteasome inhibitor MG132. The MG132 treatment abolished pIX degradation in Stop^M^-infected cells after 4 h CHX co-treatment (Fig. 4C), suggesting that pIX was degraded in a proteasome-dependent manner. Despite some experimental variability, MG132 treatment significantly stabilized pIX in Stop^M^-infected cells (Fig. 4D). To further test whether the ubiquitin-proteasome system (UPS) regulates pIX stability, we infected H1299 cells expressing His-tagged ubiquitin (His-Ubi) with Wt or Stop^M^, followed by Ni-NTA affinity purification of the His-Ubi-containing proteins. Multiple ubiquitinated pIX species (1×Ubi to 3×Ubi) were detected in Wt-infected cells, suggesting that pIX is indeed targeted by the UPS during infection (Fig. S2). In canonical proteasomal degradation, ubiquitin is attached to lysine (K) residues on target proteins (32, 33). HAdV-C5 pIX contains two lysine residues, K99 and K132 (Fig. 4E and Fig. S3A). Hence, we hypothesized that mutating K99 and K132 to arginine (R) would generate a more stable, ubiquitination-resistant pIX. To test this, we transiently expressed pIX(wt) or the K99R/K132R double mutant (referred to as pIX(2K>R)), along with His-Ubi, and purified ubiquitinated proteins. Similar to virus infections (Fig. S2), pIX(wt) displayed robust ubiquitination, as evidenced by four discrete His-Ubi-containing species (Fig. 4F). Because total pIX(2K>R) levels were also lower compared to pIX(wt) (Fig. 4F, lanes 2 to 5), the apparent decrease in ubiquitination likely reflects reduced protein abundance rather than a loss of His-Ubi conjugation. Transient expression experiments do not necessarily correlate with the data obtained from virus-infected cells. Therefore, we generated Wt and Stop^M^ viruses expressing pIX(2K>R) (hereafter as Wt(2K>R) and Stop^M^(2K>R)), assuming that these mutations may increase pIX levels in infected cells. Contrary to this prediction, and consistent with our transient transfection experiment (Fig. 4F), both Wt(2K>R) and Stop^M^(2K>R) exhibited drastically reduced expression of pIX as well as early (Flag-E1B19K, E1B55K) and late (capsid) viral proteins (Fig. 4G). The 2K>R mutation, in both plasmid and virus contexts, likely introduces additional protein defects, thereby complicating validation of pIX ubiquitination at residues K99 and K132. However, the transient transfection experiments support the conclusion that pIX is likely ubiquitinated in a lysine-independent manner.

**Fig 4.**
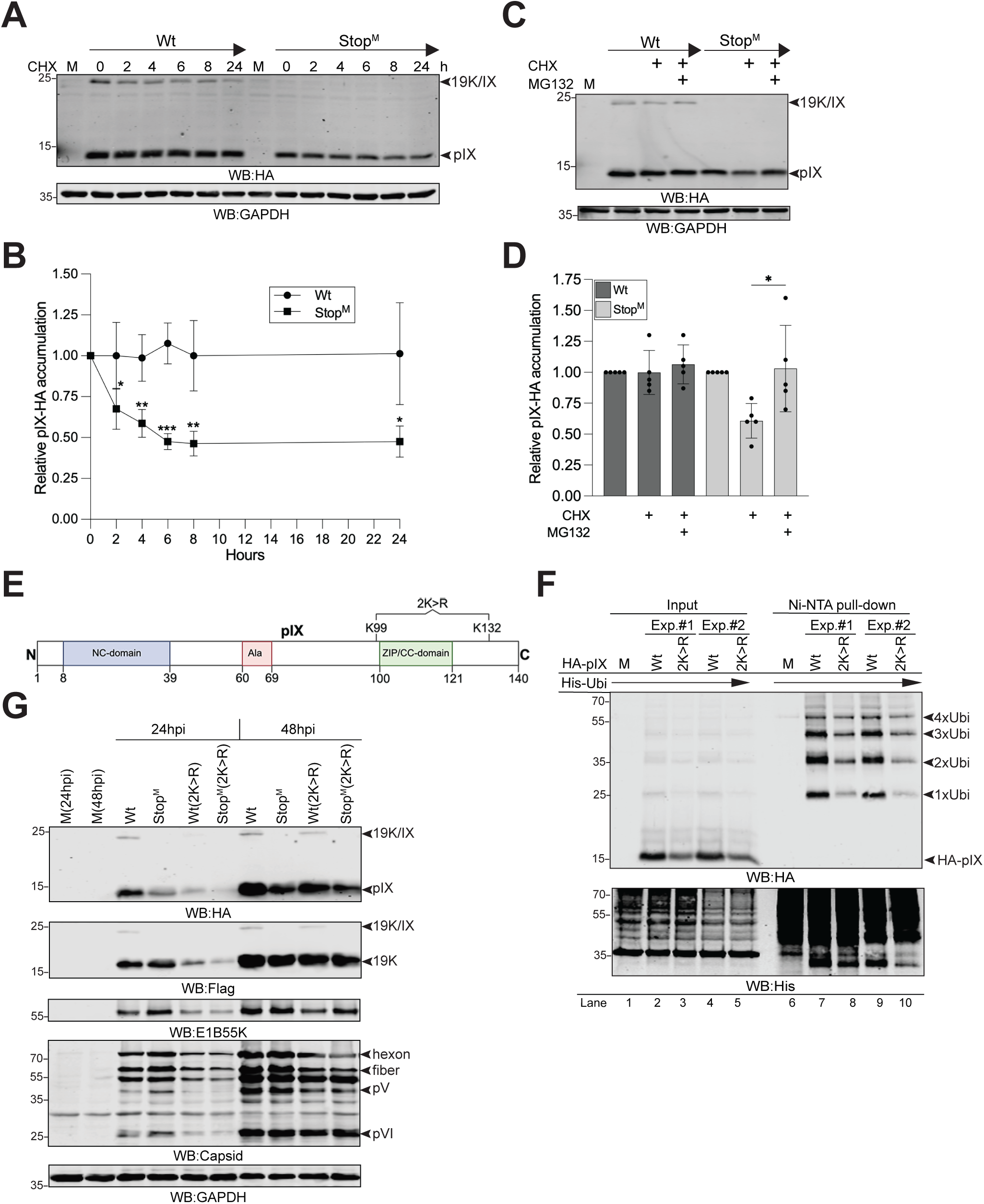
19K/IX counteracts pIX degradation in a proteasome-dependent, lysine-independent manner. **(A)** 19K/IX controls pIX stability. Wt- or Stop^M^-infected A549 cells were treated with cycloheximide (CHX) for the indicated times. A representative WB is shown. **(B)** Relative quantification of pIX-HA after normalization to GAPDH from three independent experiments (n = 3). **(C)** Proteasomal degradation of pIX. Wt- and Stop^M^-infected A549 cells were treated with CHX for 4 h in the presence or absence of the proteasome inhibitor MG132 (25 μM). **(D)** Relative quantification of independent experiments from panel C (n = 5). Quantification was performed as in panel B. **(E)** A schematic overview of the HAdV-C5 pIX with the indicated amino acids K99 and K132. The double K99R and K132R mutant is named 2K>R. **(F)** Ubiquitination of pIX outside of the infection context. His-Ubi-conjugated proteins were isolated with Ni-NTA pull-down from H1299 cells expressing His-Ubi and HA- pIX(wt) or HA-pIX(2K>R) proteins. Data from two independent experiments (Exp. #1 and #2) are shown. Distinct His-Ubi conjugates (1×Ubi to 4×Ubi) are indicated based on their migration, considering the molecular weights of pIX and ubiquitin. **(G)** 2K>R mutation does not stabilize pIX. WCL from Wt-, Stop^M^-, Wt(2K>R)-, or Stop^M^(2K>R)- infected A549 cells at 24 and 48 hpi were analyzed by WB using the indicated antibodies. M; mock, non-infected cell lysates.

### pIX is ubiquitinated at cohnserved tyrosine residues

Although ubiquitination was historically thought to occur primarily on lysine residues, subsequent studies have shown that ubiquitin can also be conjugated to cysteine (C), serine (S), threonine (T), and the N-terminus of proteins (34). Furthermore, a very recent study suggests that tyrosine (Y) residues can be ubiquitinated (35). Because HAdV-C5 pIX lacks cysteine residues (Fig. S3A), we examined whether serine or threonine residues may serve as acceptors for ubiquitin. For that purpose, we generated a set of pIX mutants in which the serine and threonine residues were mutated to alanine (referred to as S/T>A mutants) (Fig. S3B). We further immunopurified pIX, rather than using Ni-NTA purification of His-Ubi-conjugated proteins, to enrich for pIX. All immunopurified V5-tagged pIX S/T>A mutant proteins were ubiquitinated in the presence of co-expressed HA-ubiquitin (HA-Ubi) (Fig. S3C), suggesting that pIX ubiquitination does not occur at serine or threonine residues. Ubiquitination of pIX is not due to the presence of the V5 antibody epitope tag, as an irrelevant V5-tagged virus protein, pVII(K26R/K27R) (31), did not show detectable HA-Ubi conjugates (Fig. S3D). Notably, pIX contains two conserved tyrosine residues (Y14 and Y49 in HAdV-C5) (Fig. 5A and S3A). Mutation of both residues to alanine (referred to as 2Y>A) drastically reduced HA-Ubi conjugation to pIX (Figs. 5B and 5C). Non-canonical ubiquitin conjugation to tyrosine residues occurs through an oxyester bond, whereas canonical ubiquitination involves attachment of ubiquitin to lysine residues via an isopeptide bond. Hydroxylamine (NH_2_OH) can be used to demonstrate the formation of oxyester bonds, as it selectively cleaves oxyester linkages while leaving the isopeptide bonds intact (36). Indeed, NH_2_OH treatment of immunopurified V5-pIX-HA-Ubi conjugates resulted in a marked reduction of the HA-Ubi signal while leaving total V5-pIX protein levels unchanged (Fig. 5D). Tyrosine-specific ubiquitination was further detected on pIX-HA in the presence of V5-ubiquitin (Fig. S3E), confirming that the presence of different antibody epitopes does not interfere with the ubiquitination process. Lysine 48 (K48)-linked polyubiquitin chains are the canonical signal for 26S proteasomal degradation, whereas lysine 11 (K11)-linked polyubiquitin chains mediate proteasomal degradation of specific subsets of proteins, such as cell-cycle regulators (32). The HA-Ubi mutant K48R showed drastically reduced polyubiquitin chain formation on pIX, whereas the K11R mutant had no detectable effect (Fig. 5E). This suggests that pIX is predominantly modified with K48-linked polyubiquitin chains, the canonical signal for UPS-mediated degradation. Among the two conserved tyrosine residues, mutation of Y49 alone was sufficient to reduce pIX ubiquitination (Fig. 5F). Since the 19K/IX protein contains the full-length pIX sequence and therefore also the conserved tyrosine residues, we tested whether the 2Y>A mutation in the 19K/IX context affected protein ubiquitination. Similar to pIX, ubiquitin moieties were attached to 19K/IX, and the 2Y>A mutation reduced its ubiquitination (Fig. 5F). To test whether tyrosine mutations affect pIX subcellular localization, we examined individual (Y14A, Y49A) and double (2Y>A) pIX mutants by immunofluorescence. Expression of single point mutants (Y14A, Y49A) resulted in the formation of nuclear inclusions, although less frequently than in pIX(Wt)-expressing cells (Fig. 5G). In contrast, pIX(2Y>A) did not show any nuclear inclusions. These data suggest that the integrity of both Y14 and Y49 is required for optimal pIX ubiquitination and the formation of nuclear inclusions. To determine whether tyrosine ubiquitination reduces pIX stability, a CHX chase was performed in A549 cells transfected with plasmids expressing pIX(Wt) or pIX(2Y>A). The pIX(2Y>A) mutant exhibited increased resistance to CHX treatment, supporting the role of residues Y14 and Y49 in controlling pIX stability (Figs. 5H and 5I).

**Fig 5.**
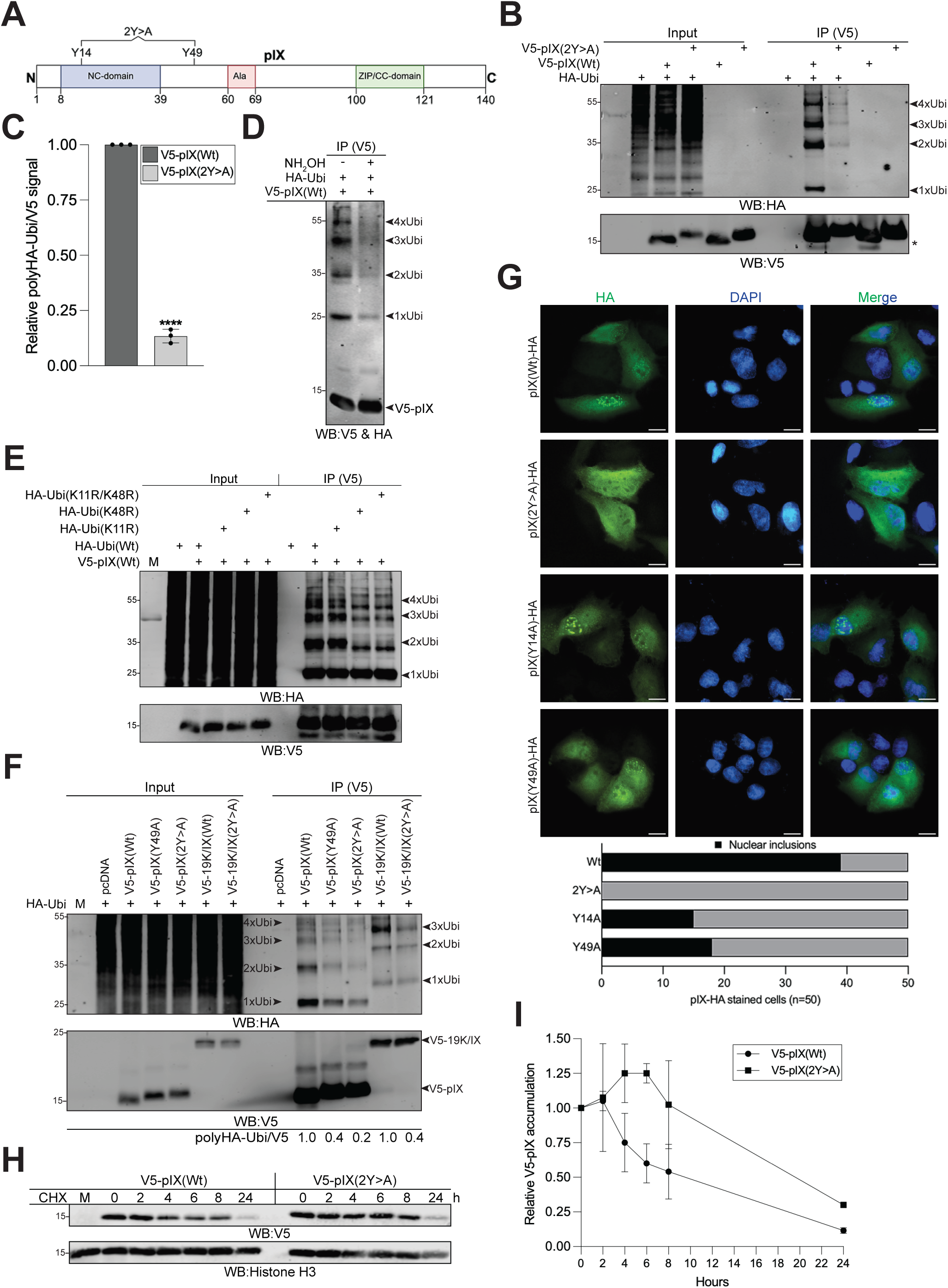
pIX is ubiquitinated at conserved tyrosine residues. **(A)** A schematic overview of pIX with the indicated tyrosine residues Y14 and Y49. The double Y14A and Y49A mutant is named 2Y>A. **(B)** Non-canonical ubiquitination of pIX. V5-pIX proteins (Wt and 2Y>A) were isolated from H1299 cells transiently expressing the HA-ubiquitin (HA-Ubi). V5-tagged pIX was immunoprecipitated (IP) with the V5 antibody, followed by protein detection by WB with the anti-V5 and anti-HA antibodies. Distinct HA-Ubi conjugates (1×Ubi to 4×Ubi) are indicated. A representative WB is shown. **(C)** Relative quantification of immunopurified polyHA-Ubi (1×Ubi to 4×Ubi) signal after normalization to the V5 IP signal from independent experiments (n = 3), shown in panel B. **(D)** Hydroxylamide (NH_2_OH) treatment blocks V5-pIX ubiquitination. V5 IP of V5-pIX/HA-Ubi conjugates was treated with NH_2_OH to break oxyester bonds between tyrosine and HA-Ubi. WB stained with the anti-V5 (detects V5-pIX) and anti-HA (detects HA-Ubi conjugates) antibodies. **(E)** V5-pIX was isolated from H1299 cells transiently expressing HA-Ubi(Wt), (K11R), (K48R), or (K11R/K48R) as in panel B. **(F)** V5-pIX and V5-19K/IX proteins (Wt, Y49A, and 2Y>A) were isolated from H1299 cells transiently expressing HA-Ubi. Protein analysis as in panel B. Distinct HA-Ubi conjugates are indicated for V5-19K/IX (1×Ubi to 3×Ubi, right side of the image) and V5-pIX (1×Ubi to 4×Ubi, left side of the image). Relative quantification (n =1) is shown below the image and was done as in panel C. **(G)** The 2Y>A mutant failed to form nuclear inclusions. Immunofluorescence in A549 cells transfected with plasmids expressing pIX(wt)-HA, pIX(Y14A)-HA, pIX(Y49A)-HA or pIX(2Y>A)-HA. Proteins were stained with the HA antibody, and nuclei were stained with DAPI at 24 h post-transfection. Scale bar = 10µm. **(H)** Increased stability of the V5-pIX(2Y>A) protein. A549 cells were transfected with plasmids expressing V5-pIX(Wt) or V5-pIX(2Y>A) for 24 h, then treated with CHX for the indicated hours (h). **(I)** Relative quantification of V5-pIX levels, normalized to histone H3, from panel H. Data are from two independent experiments (n = 2).

To understand the biological consequences of Y14 and Y49 ubiquitination during virus infection, we generated viruses expressing pIX(2Y>A) in Wt and Stop^M^ backgrounds (referred to as Wt(2Y>A) and Stop^M^(2Y>A)). Based on transient transfection experiments, we anticipated that the 2Y>A mutation would stabilize pIX in virus-infected cells. Surprisingly, the 2Y>A mutation markedly reduced pIX and capsid protein (pV and fiber) expression at 24 hpi and, to some extent, at 48 hpi (Fig. 6A). A corresponding reduction in pIX mRNA levels was also observed (Fig. 6B). Since pIX stabilizes the viral particle, the pIX(2Y>A) mutation may affect virus uptake and infectivity. However, expression of the early viral protein E1A was not reduced, indicating that the Wt and 2Y>A viruses infected cells with similar efficiency (Figs. 6A and 6B). Because the 2Y>A viruses failed to induce nuclear inclusions (Fig. 6C), the absence of these subnuclear structures may affect pIX mRNA levels. Further, given its reported transcription-activating activity (22–24), pIX might stimulate its own expression, whereas the 2Y>A mutant could be defective in this function. However, neither pIX(Wt) nor pIX(2Y>A) affected transcription from the pIX promoter in reporter assays (Fig. S4). Thus, the 2Y>A mutations may instead affect pIX transcript stability. The morphology of viral replication centers was similar in cells infected with either the Wt or 2Y>A virus (Fig. 6D), indicating that neither pIX(Wt) nor pIX(2Y>A) affects viral DNA-binding protein-positive (DBP^+^) compartments. Also, the pIX(Wt) nuclear inclusions did not localize to the DBP^+^ compartments, consistent with earlier findings (37).

**Fig 6.**
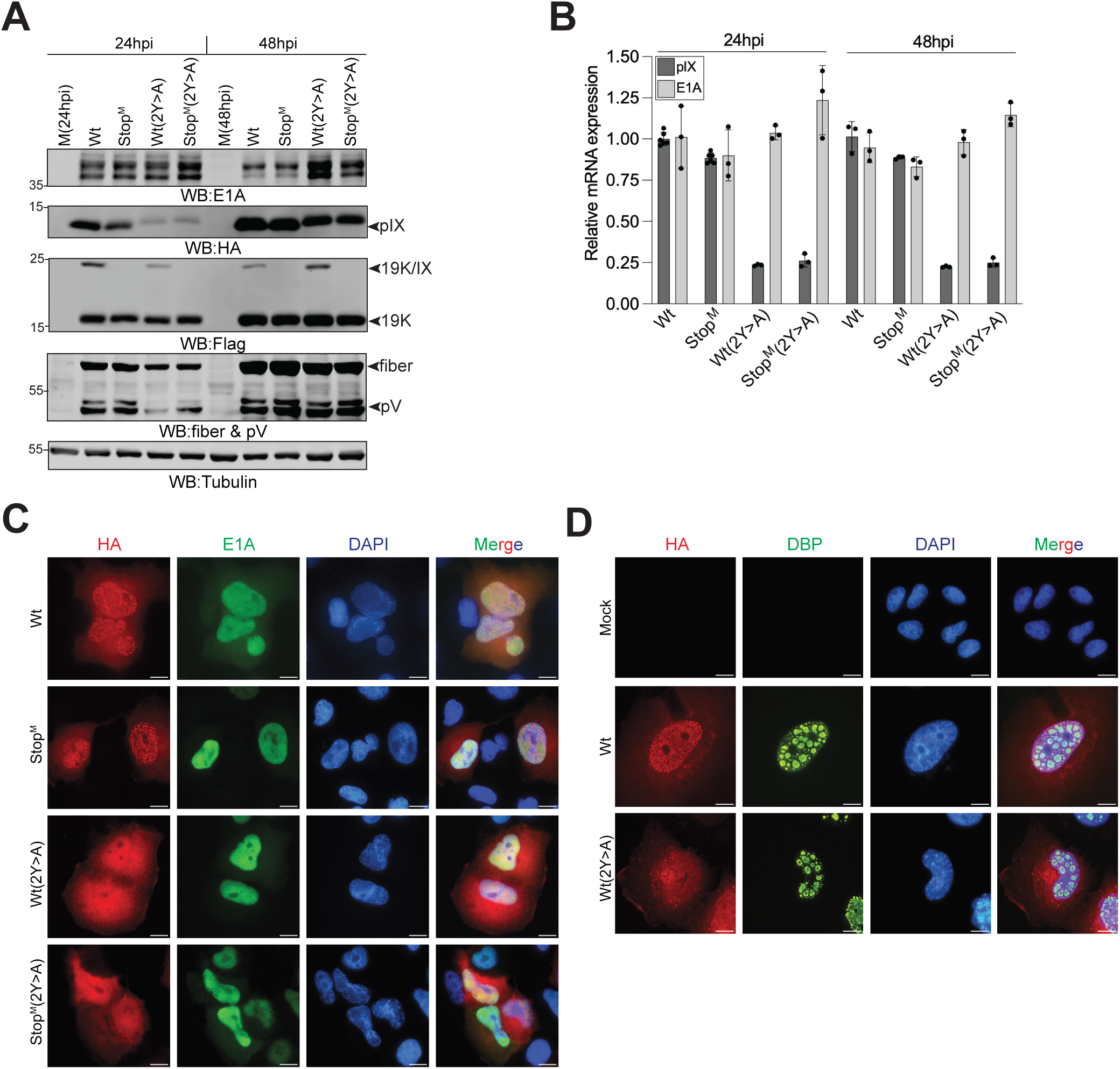
2Y>A mutation reduces pIX expression in virus-infected cells. **(A)** A549 cells were infected with Wt, Stop^M^, Wt(2Y>A), or Stop^M^(2Y>A) viruses for 24 and 48 hpi. WCLs were analyzed by WB using the indicated antibodies. E1A and capsid proteins (pV and fiber) were used as early and late phase infection markers, respectively. **(B)** Virus mRNA (pIX and E1A) expression in A549 cells infected as in panel A. Relative mRNA expression is shown after data normalization to HRPT1 expression and considering Wt-infected samples at 24 hpi as 1. **(C)** The 2Y>A mutant failed to form nuclear inclusions. Immunofluorescence in A549 cells infected with Wt, Stop^M^, Wt(2Y>A), or Stop^M^(2Y>A) viruses. Proteins were stained with the E1A (infection early phase marker) and HA antibodies, and nuclei were stained with DAPI at 24 hpi. Scale bar = 10µm. **(D)** Wt and Wt(2Y>A) do not alter viral replication centers. IF in A549 cells infected with Wt or Wt(2Y>A) viruses. Proteins were stained with the DBP (viral replication center marker) and HA antibodies, and nuclei were stained with DAPI at 24 hpi. Scale bar = 10µm.

The 2Y>A mutation likely introduces additional defects in pIX, complicating validation of ubiquitination at Y14 and Y49 in virus-infected cells. Taken together, these data indicate that pIX undergoes non-canonical tyrosine ubiquitination, thereby reducing its stability in transiently transfected cells.

### pIX interacts with specific 26S proteasome subunits

To further investigate whether ubiquitination of pIX targets the protein for degradation, pIX(wt)-HA was immunopurified from Stop^M^-infected A549 cells. Interacting proteins were identified by mass spectrometry (MS)-based proteomics, and enrichment was compared between non-epitope-tagged HAdV-C5 (Wt_AB) and HA-tagged Stop^M^ virus samples. As expected, pIX interacted with its known partner, the hexon protein (Fig. 7A). In addition, several HAdV-C5 proteins, pVIII, IVa2, fiber, E1B-55K, pIIIa, and DBP were identified as novel pIX binding partners (Fig. 7A). Interestingly, the most prominent cellular pIX interactor was the 26S proteasome subunit PSMC3 (Rpt5), which, together with another pIX-binding protein, PSMD9 (p27, Rpn4) (Fig. 7A), forms the p27 precursor complex needed for 26S proteasome assembly (38). To validate the MS findings, we repeated the experiment in Wt- and Stop^M^-infected A549 cells, confirming specific interactions between pIX-HA and PSMC3 and PSMD9 (Fig. 7B). In contrast, the unrelated adenovirus protease (AVP) showed no binding, confirming interaction specificity. Because pIX-HA levels were reduced in Stop^M^-infected cells (Fig. 2B), quantitative IP analysis normalized to pIX-HA revealed enhanced pIX-HA binding to both proteins in the absence of 19K/IX (Fig. 7B). Immunofluorescence analysis further confirmed that both PSMC3 and PSMD9 colocalize with pIX-HA in the cytoplasm, while PSMD9 may additionally localize with pIX in the nucleus (Fig. 7C). Since ubiquitin can be conjugated to the Y14 and Y49 residues (Fig. 5B), we tested whether ubiquitination-deficient pIX(2Y>A) has altered binding to PSMC3 and PSMD9. Notably, pIX(2Y>A) showed reduced binding to PSMC3, whereas binding to PSMD9 remained essentially unchanged in transiently transfected cells (Fig. 7D). To further investigate the impact of pIX-PSMC3/PSMD9 interactions, PSMC3 and PSMD9 were overexpressed in Wt- and Stop^M^-infected cells, and pIX expression was analyzed at 24 hpi. Overexpression of both proteins reduced pIX accumulation more severely in Stop^M^-infected cells, suggesting that the absence of 19K/IX enhances pIX degradation in a PSMC3- and PSMD9-dependent manner (Fig. 7E, lanes 6 to 9 and quantification in Fig. 7F).

**Fig 7.**
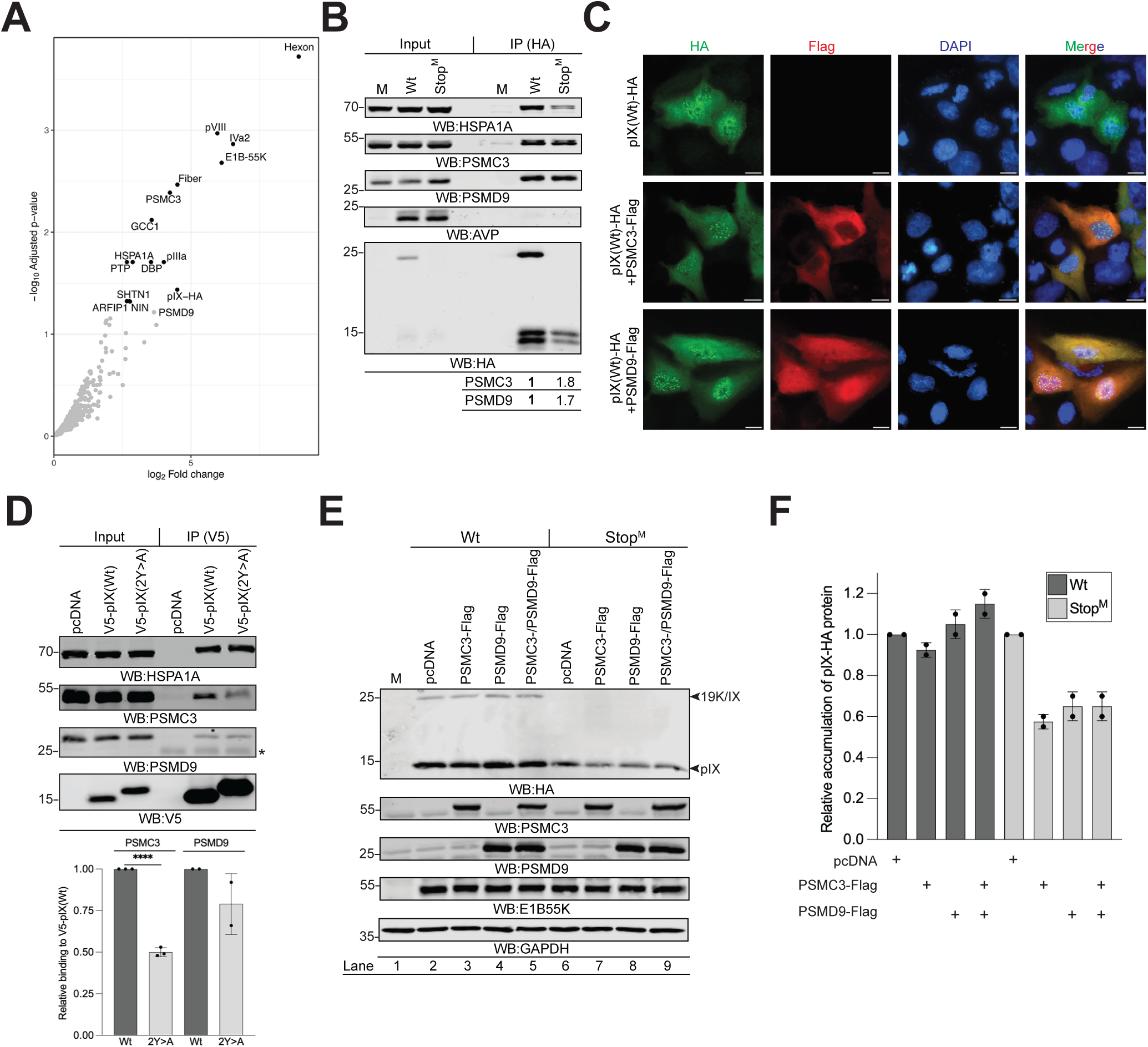
pIX interacts with the 26S proteasome subunits. **(A)** pIX-HA interacts with the 26S proteasome subunits. Volcano plot of the MS experiment showing identified proteins that interact with pIX-HA in Stop^M^ (MOI=10) infected A549 cells. An adjusted *P*-value cut-off of 0.05 and a log_2_ fold change cut-off of 2 were used. Data are shown from a biological triplicate experiment (n=3). **(B)** Validation of pIX-HA interacting proteins. The pIX-HA was immunopurified from A549 cells infected with Wt or Stop^M^ at 48 hpi. WB analysis with antibodies against HSPA1A, PSMC3, PSMD9, HA, and AVP (adenoviral proteinase, used as a negative control). Relative binding of the PSMC3 and PSMD9 proteins to the immunopurified pIX-HA is shown. **(C)** Immunofluorescence in A549 cells transfected with the plasmids expressing the indicated proteins at 24 hpt. Proteins were stained with the HA and Flag antibodies, and nuclei were stained with DAPI. Scale bar = 10 µm. **(D)** Tyrosine mutation (2Y>A) reduces pIX binding to the 26S proteasome subunit PSMC3. WB analysis of V5-pIX complexes with the indicated antibodies. Relative binding of PSMC3 and PSMD9, normalized to immunopurified V5-pIX(Wt) and V5-pIX(2Y>A) from three independent experiments (n=3), is shown below the image. **(E)** Overexpression of Flag-PSMC3 and Flag-PSMD9 reduces pIX accumulation in virus-infected cells. Wt or Stop^M^-infected A549 cells were also transfected with plasmids expressing Flag-tagged PSMC3 or PSMD9 for 24 h. WCLs were analyzed by WB with the indicated antibodies. **(F)** Relative accumulation of pIX-HA protein normalized to GAPDH, from two independent experiments shown in panel E (n = 2).

## Discussion

Recent long-read transcriptomic analyses have revealed an extraordinary level of complexity of the HAdV-C2 and HAdV-C5 transcriptomes (12–14). These studies identified unique transcripts in which alternative splicing links CDSs from different viral transcription units, potentially generating novel in-frame fusion proteins. Given that the 19K/IX protein comprises one-third of the E1B19K CDS and the entire pIX CDS, it may exhibit biochemical characteristics of both proteins. Based on our biochemical fractionation and IF experiments, the 19K/IX protein shows a subcellular localization pattern similar to pIX in infected cells. Although 19K/IX contains the BH3 motif of E1B19K, neither our DSS cross-linking experiments nor MS analysis provided clear evidence for the E1B19K-pIX interaction. The observation that both proteins, pIX and 19K/IX, exhibit overlapping subcellular localization implies that these two proteins function in a protein complex. Our protein-protein interaction and DSS cross-linking studies confirmed this, revealing pIX-19K/IX heterodimers in both virus-infected and transfected cells. Biochemical studies have shown that the C-terminal ZIP/CC-domain mediates pIX self-assembly/oligomerization (24, 29). Furthermore, several structural studies have shown that pIX forms trimeric structures that bind the hexon protein and stabilize the virion structure (20, 39). It is therefore possible that pIX, which accumulates at higher levels than 19K/IX, could form a trimeric pIX-19K/IX-pIX complex. However, because our DSS cross-linking experiments did not reveal such a trimeric complex in infected cells, we favor a model in which a heterodimeric pIX-19K/IX complex is formed during infection (Fig. 8).

**Fig 8.**
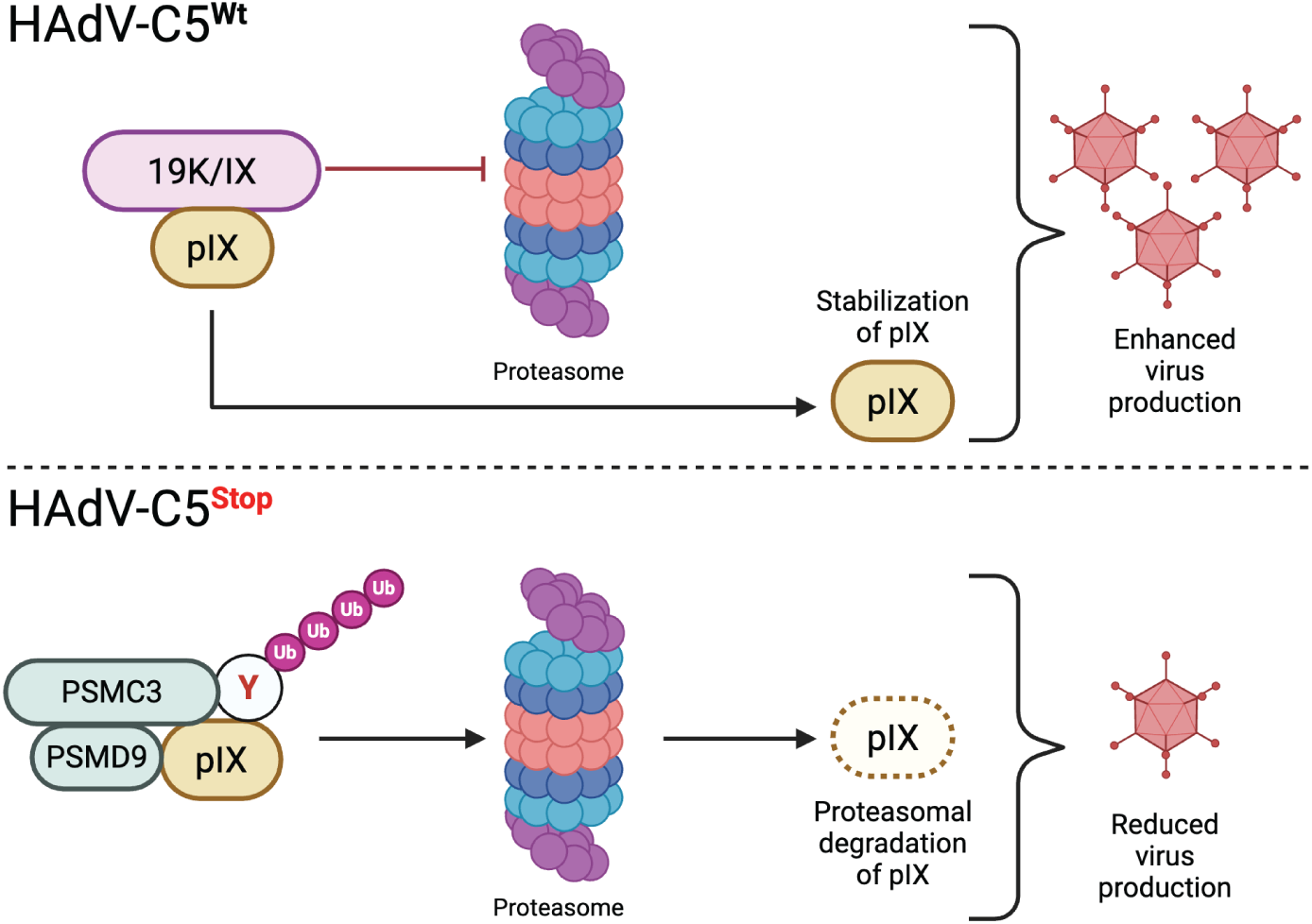
Proposed model of 19K/IX interplay with the pIX in HAdV-C5 life cycle. The binding of 19K/IX to pIX in HAdV-C5(Wt)-infected cells blocks pIX proteasomal degradation and thereby enhances infectious virus progeny production. Lack of 19K/IX and pIX ubiquitination at residues Y14 and Y49 mediates binding of PSMC3 and PSMD9 to pIX, followed by proteasomal degradation of pIX.

Mechanistically, we show that 19K/IX stabilizes pIX by blocking its proteasomal degradation during infection (Fig. 8). A striking finding of our study is the identification of non-canonical tyrosine ubiquitination of pIX. Since ubiquitin conjugation via an oxyester bond is considered chemically labile (40), detecting tyrosine ubiquitination has remained enigmatic. Consequently, the prevalence and biological relevance of tyrosine ubiquitination may be underestimated. A recent study provides the first evidence that tyrosine ubiquitination can occur *in vivo* (35). Our study builds on this work by identifying pIX as an additional protein that can be ubiquitinated on tyrosine residues. Protein ubiquitination does not necessarily lead to protein degradation, as the functional outcome depends on the type of ubiquitin linkage and the site of ubiquitin conjugation on the target protein (41, 42). In this regard, pIX ubiquitination at tyrosine residues appears to follow the classical UPS, whereby ubiquitinated proteins are targeted for proteasomal degradation. This is supported by our observations that pIX undergoes K48-linked polyubiquitination and that MG132 treatment stabilized pIX in virus-infected cells. Furthermore, pIX interacted specifically with the proteasome subunits PSMC3 and PSMD9, consistent with ubiquitinated pIX being targeted for proteasomal degradation. The 26S proteasome consists of the 19S regulatory particle (19S RP), which recognizes ubiquitinated substrates, and the 20S core particle (20S CP), which has the proteolytic activity (43). Proteasomal degradation is initiated by docking of ubiquitinated proteins to ubiquitin receptors on the 19S RP. Notably, PSMC3 and PSMD9 are not ubiquitin receptors. Rather, PSMC3 is an ATPase subunit of the 19S RP, and PSMD9 is a 19S RP assembly chaperon (43, 44). PSMD9 and PSMC3, along with the third subunit, PSMC6, form a stable subcomplex, known as the p27 precursor complex, that facilitates assembly of the 19S RP (38, 45, 46). Because PSMD9 functions as a chaperone and is not present in the mature 26S proteasome (38), we propose that the p27 precursor complex interacts with pIX via Y14 and Y49 through the PSMC3 subunit. Thereafter, pIX is delivered to the 26S proteasome via the p27 precursor complex, where it undergoes proteasomal degradation. Thus, 19K/IX likely functions to reduce pIX delivery to the 26S proteasome via PSMC3 and PSMD9 (Fig. 8).

In summary, our study provides functional insights into a novel HAdV-C5 in-frame fusion protein, 19K/IX. Notably, E1-deleted HAdV vectors are widely used as gene delivery methods and do not express the 19K/IX protein. Given the stabilizing effect of 19K/IX on pIX, this finding raises the possibility that 19K/IX expression in producer cells could influence HAdV vector production. An interesting approach for future studies could be to determine whether 19K/IX expression can enhance HAdV vector quality and yields.

## Materials and methods

### Cell lines

The A549, H1299, and HEK293AD cells were obtained from ATCC and Cell Biolabs. All cell lines were grown in DMEM (Thermo Fisher Scientific) supplemented with 10% fetal bovine serum (FBS, Thermo Fisher Scientific) and penicillin-streptomycin solution (PEST; Thermo Fisher Scientific) at +37°C in a 5% CO_2_ incubator.

### Plasmids and transfections

Detailed descriptions of the plasmids are available in the Supplementary Materials and Methods, and in Table S1. All plasmid transfections were performed with the jetPRIME (Polyplus) transfection reagent (30).

### Construction of HAdV-C5 mutants and virus infections

Replication-competent HAdV-C5 viruses (Wt, Stop^M,^ or 5’ss^M^) were generated using the AdenoBuilder system (47, 48). Detailed virus construction and isolation approaches are described in the Supplementary Materials and Methods. Unless otherwise indicated in the figure legends, virus infections were performed at a multiplicity of infection (MOI) of 5, defined as fluorescence-forming units (FFU/cell), in infection medium (DMEM with 2% Newborn Calf Serum and PEST) for 1 h at +37°C, as described previously (30). All used viruses are described in Table S2.

### Virus titration

All viruses were titered simultaneously in A549 cells using an anti-hexon antibody (TC31-9C12.C9-s, DSHB; Table S3). The detailed protocol is available in the Supplementary Materials and Methods.

### RNA isolation and transcript detection by qRT-PCR and PCR

Total RNA was extracted with TRIreagent (Sigma, T9424) according to the manufacturer’s protocol. cDNA synthesis and qRT-PCR are described in the Supplementary Materials and Methods. Oligonucleotide sequences are described in Table S4. Endpoint PCR reactions were performed with PrimeSTAR HS DNA polymerase (Takara) using primers that anneal to the 5’ and 3’ ends of the Flag-19K/IX-HA sequence (Table S4). PCR products were analyzed on a 1% agarose gel in 1× TAE buffer, gel-purified, and Sanger-sequenced (Eurofins) with the same oligos used for PCR amplification. DNA sequences were analyzed with CLC Main Workbench (Qiagen) using the updated HAdV-C5 transcript database described in (14).

### DSS crosslinking

The disuccinimidyl suberate (DSS, Thermo Fisher Scientific, A39267) was initially dissolved in DMSO (50 mM) and was further diluted to final concentrations of 0.1, 0.3, and 1 mM in PBS. Before crosslinking, infected A549 cells were washed once with PBS, then incubated in the PBS/DSS solution for 30 min at room temperature. The reaction was quenched with 20 mM Tris-HCl (pH 7.5) for 15 min at room temperature. After quenching, the cells were washed twice with PBS and lysed in RIPA buffer (50 mM Tris pH 7.5, 150 mM NaCl, 1% NP40, 1% sodium deoxycholate, 0.1% sodium dodecyl sulfate) for protein analysis.

### Cycloheximide chase

Virus-infected or plasmid-transfected cells were treated with cycloheximide (CHX, Sigma, C4859, 100 μg/ml as final concentration) 24 hours post-transfection (hpt) or 24 hpi, respectively. Treated cells were collected at 2, 4, 6, 8, and 24 h after CHX was added to the growth medium, lysed in RIPA buffer, and analyzed by Western blotting (WB) as described previously (49). Relative pIX intensity was calculated after normalization to the GAPDH or histone H3 proteins. The protein signal at the time of adding CHX (t=0) was considered as 1. CHX and MG132 co-treatments were performed by incubating cells simultaneously with CHX (100 µg/ml) and MG132 (25 µM) for 4 h. DMSO was used as the vehicle control.

### Immunoprecipitation

Immunoprecipitation (IP) was performed from A549 or H1299 cells transfected with plasmids or infected with viruses. At 24 hpt or 24 hpi, the cells were lysed with Pierce IP lysis buffer (Thermo Fisher Scientific, #87788) supplemented with protease inhibitors (Halt Protease Inhibitor Cocktail, Thermo Fisher Scientific, #78437). The Flag-pIX protein was immunopurified from soluble cell extracts using anti-Flag (M2) magnetic beads (Sigma, M8823) at +4°C overnight. The pIX-HA and V5-pIX proteins were immunoprecipitated with HA-Trap and V5-Trap magnetic agarose (Proteintech), respectively. Beads were washed 4times in 1 ml of lysis buffer and analyzed by 13.5% SDS-PAGE. The antibodies used are described in Table S3.

### Ubiquitination assays

His-ubiquitin (His-Ubi) assays were done as described previously (50) with minor modifications. H1299 cells were infected with viruses (MOI=50), followed by His-Ubi-expressing plasmid transfection 1 hpi. At 24 hpi, the cells were treated with MG132 (25 µM) for an additional 4 h. The cells were washed with ice-cold PBS supplemented with N-ethylmaleimide (NEM, Sigma, E3876, 40 mM final concentration), collected in PBS+NEM, and finally lysed in lysis buffer. His-Ubi-conjugated proteins were isolated using Ni-NTA beads (Qiagen), and the eluted proteins were analyzed by WB. For HA-Ubi assays, cells were treated with MG132, lysed in RIPA buffer, and V5-tagged proteins were immunoprecipitated from the soluble cell lysates using V5-Trap magnetic agarose. The beads were washed with 4 × 1 ml of RIPA buffer, and the proteins were eluted with 2 × Laemmli sample buffer and analyzed by 13.5% SDS-PAGE.

### Hydroxylamine treatment

H1299 cells were transfected with plasmids expressing V5-pIX(wt) and HA-Ubi for 24 h. V5-pIX(wt) was immunopurified as above, and the V5-pIX(wt)/HA-Ubi conjugates were released from the magnetic beads with 2% SDS in PBS. The eluate was divided into two equal fractions and treated with hydroxylamine (NH_2_OH, Sigma, 467804, final concentration 1.5 M) for 1 h at 37°C. In the control reaction, NH_₂_OH was replaced with an equal volume of H_2_O, and samples were incubated in parallel with NH_₂_OH-treated reactions. The reactions were stopped with 6 × Laemmli sample buffer, heated at +95°C for 5 min, and analyzed on 13.5% SDS-PAGE.

### Subcellular fractionation

Cell fractionation was done using the Cell Fractionation Kit (Abcam, ab109719) with minor modifications described in (30). The quality of the fractions was analyzed by western blotting using antibodies against Tubulin, CoxIV, and histone H3, respectively.

### SDS-PAGE and Western blotting

In experiments in which whole-cell lysates (WCLs) were analyzed, cells were lysed in RIPA buffer with Halt inhibitors. Proteins were detected on 0.2 µm nitrocellulose membranes with primary antibodies (Table S3) and fluorescence-labeled secondary antibodies (IRDye, LI-COR) followed by visualization with the Odyssey CLX imaging system (LI-COR). Protein signals were quantified using the Image Studio (LI-COR) software.

### Mass spectrometry

Sample preparation and mass spectrometry (MS) experiments were done as previously described (49) with minor modifications. A549 cells were infected (MOI=10) in triplicate with HAdV-C5 (Wt_AB, Table S2) and Stop^M^ viruses for 48 hpi. WCLs were prepared as described in (49), and pIX-HA was immunopurified using HA-Trap magnetic beads (Proteintech). A detailed description of the MS analysis is provided in the Supplementary Materials and Methods.

### Indirect immunofluorescence assay

A549 cells were grown on coverslips in a 24-well plate and infected or transfected with the respective viruses (MOI=5) or plasmids (100 ng plasmid DNA). Cells were fixed in 4% PFA in PBS for 10 min and permeabilized with 0.1% Triton X-100 in PBST (PBS + 0.01% Tween-20) for 10 min at room temperature, as in (30). The primary antibodies used are listed in Table S3. Proteins were visualized with the secondary antibodies (Alexa Fluor 594, A21442, and Alexa Fluor 488, A11001; Thermo Fisher Scientific) along with the DAPI solution (Sigma, D9542, final conc. 300 nM) for 1 h at room temperature. The slide was mounted with ProLong Diamond Antifade Mountant (Thermo Fisher Scientific) and analyzed with a fluorescence microscope (Nikon Eclipse 90i) using NIS-Elements (Nikon) software.

### Statistical analysis

Statistical analysis was performed in GraphPad Prism (version 9.4.0) statistical software (GraphPad). Data are presented as mean ± standard deviation (SD). An unpaired *t*-test was used to assess statistical significance (****; *p* ≤ 0.0001, ***; *p* ≤ 0.001, **; *p* ≤ 0.01; *, *p* ≤ 0.05;, *p* > 0.05; ns, not significant).

## Data availability

The mass spectrometry proteomics data have been deposited to the ProteomeXchange Consortium (http://proteomecentral.proteomexchange.org) via the PRIDE partner repository (51) with the dataset identifier PXD076227.

## Acknowledgments

We thank Prof. Göran Akusjärvi and Prof. Catharina Svensson for continuous support; Dr. Jinlin Li for comments on the manuscript; Noemi Zsuzsa Kovacs and Charlotte Rex for technical assistance; Dr. David Matthews, Dr. Arnold Levin, Dr. Thomas Dobner, and Dr. Maxim Balakirev for providing antibodies; and Dr. Daniel Öberg for the initial identification of the 19K/IX transcript. Vasileios Rafail Evangelopoulos was supported by a generous fellowship from the Greek State Scholarships Foundation (IKY). The Swedish Cancer Society (180599 and 211537) and Knut and Alice Wallenberg Foundation (KAW 2017.0071) have financially supported this study.

