## Supplementary Figures,Tables & Materials for "A novel adenovirus 19K/IX protein enhances infection by attenuating proteasomal degradation of tyrosine-ubiquitinated capsid protein IX"

#### Supplementary Figure Legends

**Fig S1. Expression and subcellular localization of the pIX and 19K/IX proteins. (A)** Lack of 19K/IX reduces pIX expression. A549 cells were infected (MOI = 5) with Wt or $Stop^{M}$, and whole-cell lysate (WCL) was prepared at 12, 16, 20, and 24 hpi. WCLs were analyzed by western blotting (WB) with the anti-HA, anti-Flag, anti-E1A, anti-fiber, and anti-tubulin antibodies. **(B)** pIX induces nuclear inclusions. Immunofluorescence (IF) of A549 cells infected (MOI = 5) with Wt or $Stop^{M}$ at 24 hpi. pIX-HA was stained with the HA antibody, and nuclei were stained with DAPI. Scale bar = 10 µm. **(C)** pIX is relocalizing 19K/IX into nuclear inclusions. IF in A549 cells transiently transfected with plasmids expressing HA-pIX, V5-19K/IX, or both proteins at 24 hpt. The proteins were stained with the HA and V5 antibodies, and nuclei were stained with DAPI. Scale bar = 10 µm. **(D)** The same IF as in panel C, but performed in H1299 cells at 24 hpt. **(E)** Epitope tagging does not change 19K/IX localization. IF in A549 cells transiently transfected with plasmids expressing HA-pIX, V5-19K/IX, or both proteins at 24 hpt. The proteins were stained with the HA and V5 antibodies, and nuclei were stained with DAPI. Scale bar = 10 µm.

**Fig S2. pIX is ubiquitinated in infected cells.** H1299 cells transiently expressing His-ubiquitin (His-Ubi) were infected with Wt or $Stop^{M}$ (MOI = 50) for 24 h. Ni-NTA affinity-purified His-Ubi-conjugated proteins were separated by WB and analyzed with the indicated antibodies. Ubiquitinated pIX species are indicated; a red asterisk indicates Flag-19K/IX-HA.

**Fig S3. pIX is not ubiquitinated at serine and threonine residues. (A)** Amino acid alignment (Clustal Omega Multiple Sequence Alignment) of pIX proteins belonging to different HAdV types (HAdV-C5 (P03281), HAdV-A12 (P03284), HAdV-F41 (P32539), HAdV-G52 (A0MK45), HAdV-D37 (B2VQE5), HAdV-B7 (P68971), HAdV-E4 (Q2KSG2)). Conserved tyrosine (Y) residues (Y14 and Y49 in HAdV-C5) are boxed in red. Non-conserved lysine (K) residues (K99 and K132 in HAdV-C5) are boxed in green. **(B)** Schematic illustration of the serine (S) and threonine (T) residues in HAdV-C5 pIX. Mutations of the respective S and T residues to alanine (S/T>A) in the individual pIX proteins are shown. **(C)** Serine and threonine mutations do not change pIX ubiquitination. H1299 cells transfected with HA-Ubi and individual V5-pIX (panel B) encoding plasmids. The V5-tagged proteins were immunopurified and probed with anti-HA (green, detects ubiquitinated proteins) and anti-V5 (red, detects pIX). **(D)** V5 epitope tag (NH2-G**K**PIPNPLLGLDST-COOH) is not ubiquitinated despite the presence of the lysine (marked in bold) residue. V5-pIX(Wt) and irrelevant V5-epitope tag containing HAdV-C5 protein pVII(K26R/K27R) (referred to as V5-pVII) were immunopurified (IP) from H1299 cells transiently expressing the HA-Ubi. Proteins were detected by WB with the anti-V5 and anti-HA antibodies. Distinct HA-Ubi conjugates (1×Ubi to 4×Ubi) are indicated by their migration, based on the molecular weights of V5-pIX and HA-Ubi. **(E)** Antibody epitope tagging does not change pIX ubiquitination. V5 epitope-tagged ubiquitin (V5-Ubi) is specifically conjugated to the pIX(wt)-HA protein. Wild-type and mutant pIX-HA were immunopurified from H1299 cells transiently expressing V5-Ubi. Proteins were detected by WB with the anti-V5 and anti-HA antibodies. An asterisk indicates a potentially modified version of pIX, detected only in the case of pIX-HA.

**Fig S4. pIX does not induce gene expression from its own promoter.** A549 cells were transiently transfected with a plasmid containing the pIX promoter fused to the NanoLuc (NLuc) reporter gene together with plasmids expressing pIX(Wt)-HA or pIX(2Y>A)-HA. Relative NLuc expression was analyzed 24 h post-transfection from two independent experiments performed in technical triplicates. E1A expression was used as a positive control for pIX promoter activation.

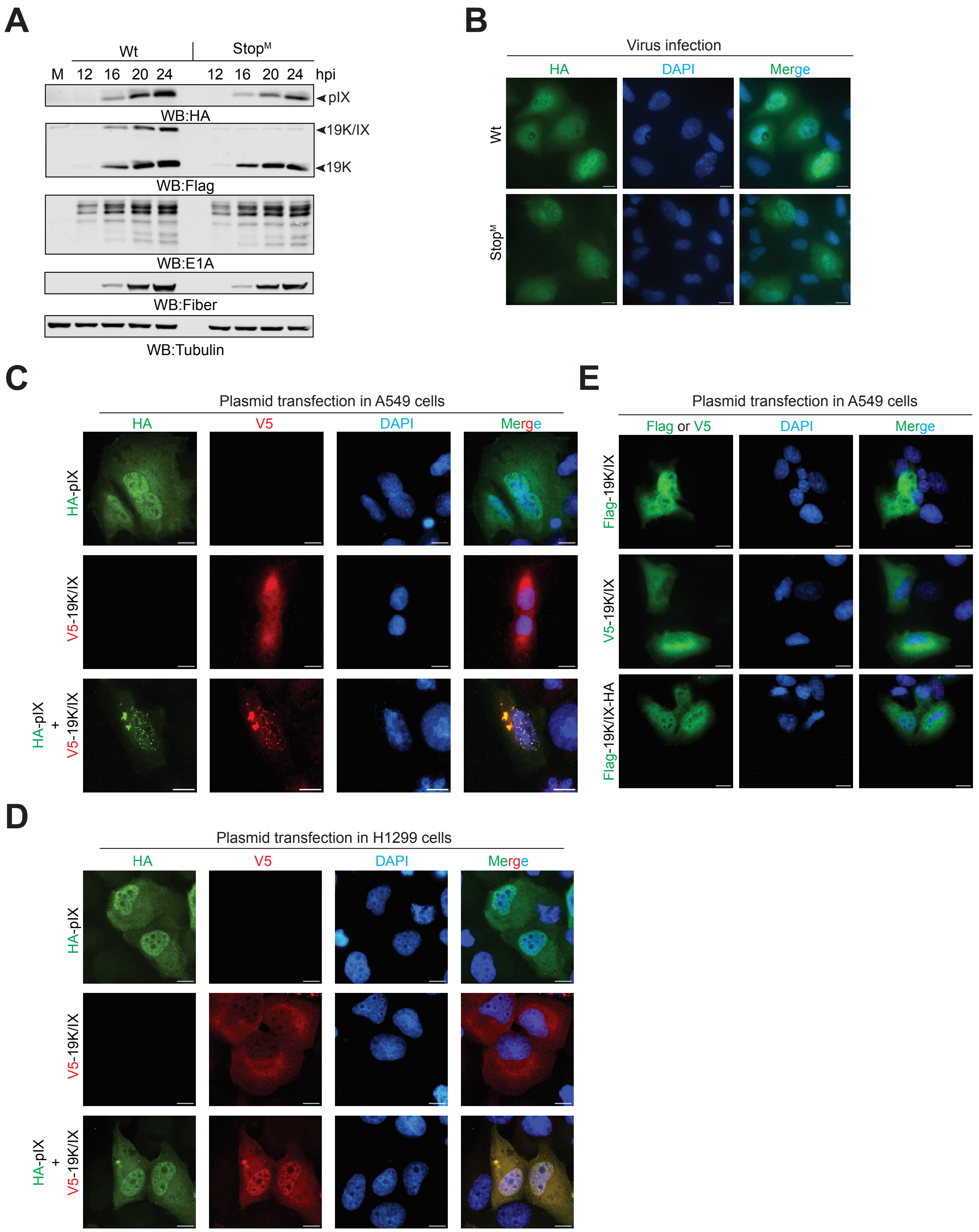
A
Wt
StopM
M 12 16 20 24 12 16 20 24 hpi
pIX
WB:HA
19K/IX
19K
WB:Flag
WB:E1A
WB:Fiber
WB:Tubulin
B
Virus infection
HA
DAPI
Merge
Wt
StopM
C
Plasmid transfection in A549 cells
HA
V5
DAPI
Merge
HA-pIX
V5-19K/IX
HA-pIX + V5-19K/IX
D
Plasmid transfection in H1299 cells
HA
V5
DAPI
Merge
HA-pIX
V5-19K/IX
HA-pIX + V5-19K/IX
E
Plasmid transfection in A549 cells
Flag or V5
DAPI
Merge
Flag-19K/IX
V5-19K/IX
Flag-19K/IX-HA

**Fig. S1**

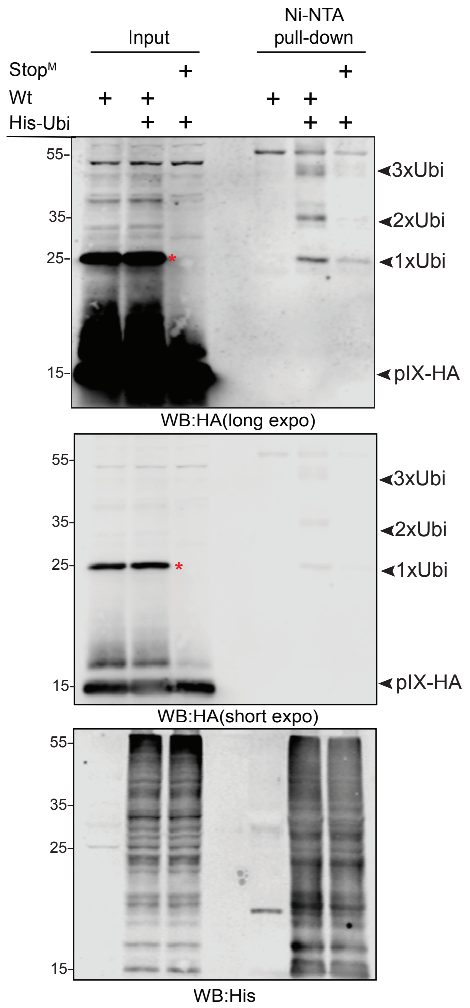
Input
Ni-NTA pull-down
Stop^M
Wt
His-Ubi
55-
35-
25-
15-
3xUbi
2xUbi
1xUbi
pIX-HA
WB:HA(long expo)
WB:HA(short expo)
WB:His

### Fig. S2

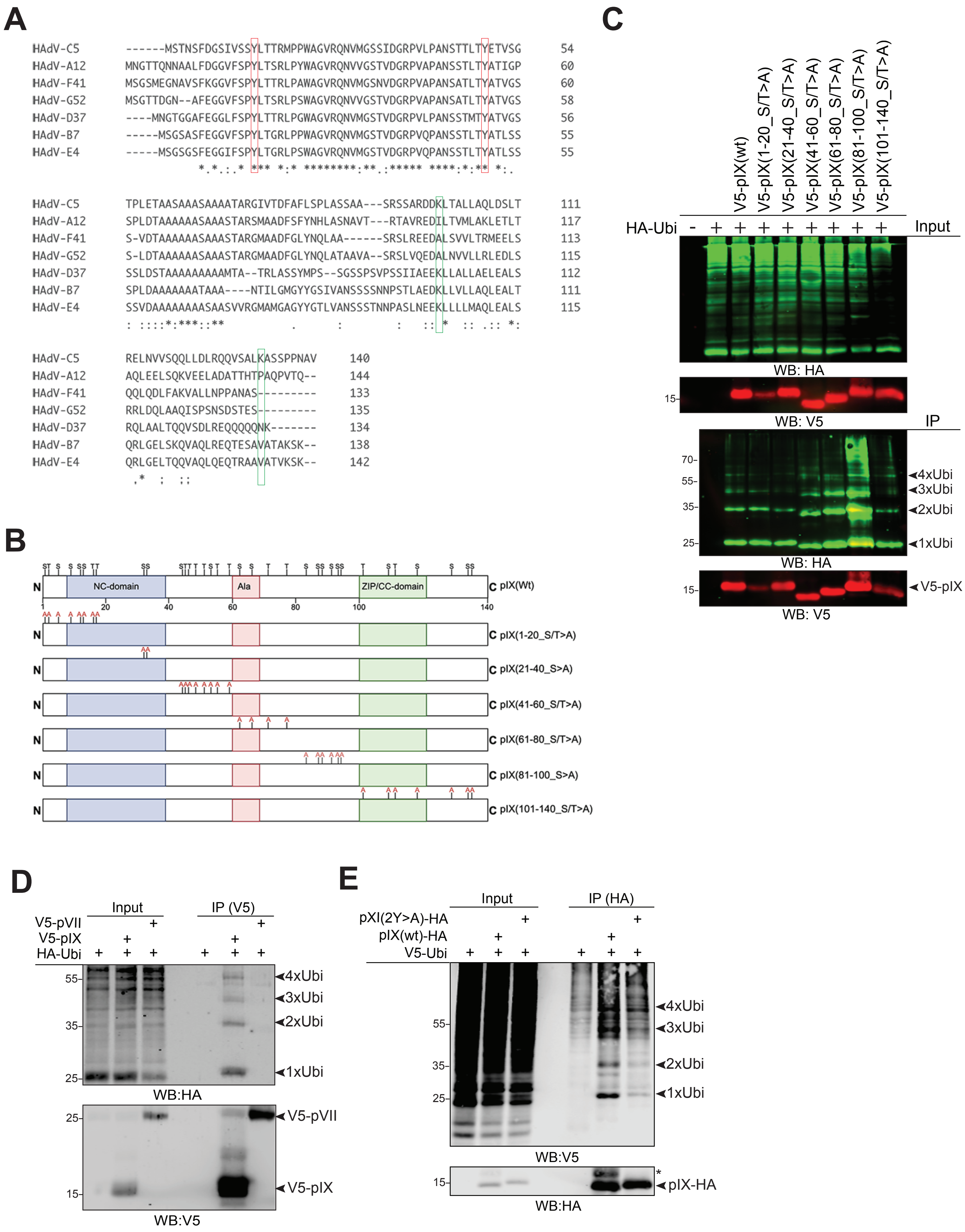
A
HAdV-C5 ------MSTNSFDGSIVSSYLTTRMPPWAGVRQNVMGSSIDGRPVLPANSTTLTYETVSG 54
HAdV-A12 MNGTTQNNAALFDGGVFSPYLTSRLPYWAGVRQNVVGSTVDGRPVAPANSSTLTYATIGP 60
HAdV-F41 MSGSMEGNAVSFKGGVFSPYLTTRLPAWAGVRQNVMGSNVDGRPVAPANSATLTYATVGS 60
HAdV-G52 MSGTTDGN--AFEGGVFSPYLTSRLPSWAGVRQNVVGSTVDGRPVAPANSATLTYATVGS 58
HAdV-D37 ----MNGTGGAFEGGLFSPYLTTRLPGWAGVRQNVMGSTVDGRPVLPANSSTMTYATVGS 56
HAdV-B7 -----MSGSASFEGGVFSPYLTGRLPPWAGVRQNVMGSTVDGRPVQPANSSTLTYATLSS 55
HAdV-E4 -----MSGSGSFEGGIFSPYLTGRLPSWAGVRQNVMGSTVDGRPVQPANSSTLTYATLSS 55
HAdV-C5 TPLETAASAAASAAAATARGIVTDFAFLSPLASSAA---SRSSARDDKLTALLAQLDSLT 111
HAdV-A12 SPLDTAAAAAASAAASTARSMAADFSFYNHLASNAVT---RTAVREDILTVMLAKLETLT 117
HAdV-F41 S-VDTAAAAAASAAASTARGMAADFGLYNQLAA------SRSLREEDALSVVLTRMEELS 113
HAdV-G52 S-LDTAAAAAASAAASTARGMAADFGLYNQLATAAVA--SRSLVQEDALNVVLLRLEDLS 115
HAdV-D37 SSLDSTAAAAAAAAAMTA--TRLASSYMPS--SGSSPSVPSSIIAEEKLLALLAELEALS 112
HAdV-B7 SPLDAAAAAAAATAAA----NTILGMGYYGSIVANSSSSNNPSTLAEDKLLVLLAQLEALT 111
HAdV-E4 SSVDAAAAAAAASAASVVRGMAMGAGYYGTLVANSSSTNNPASLNEEKLLLLMAQLEALS 115
HAdV-C5 RELNVVSQQLLDLRQQVSALKASSPPNAV 140
HAdV-A12 AQLEELSQKVEELADATTHTPAQPVTQ-- 144
HAdV-F41 QQLQDLFAKVALLNPPANAS--------- 133
HAdV-G52 RRLDQLAAQISPSNSDSTES--------- 135
HAdV-D37 RQLAALTQQVSDLREQQQQQNK------- 134
HAdV-B7 QRLGELSKQVAQLREQTESAVATAKSK-- 138
HAdV-E4 QRLGELTQQVAQLQEQTRAAVATVKSK-- 142
B
NC-domain
Ala
ZIP/CC-domain
N
C pIX(Wt)
1 20 40 60 80 100 140
C pIX(1-20_S/T>A)
C pIX(21-40_S>A)
C pIX(41-60_S/T>A)
C pIX(61-80_S/T>A)
C pIX(81-100_S>A)
C pIX(101-140_S/T>A)
C
V5-pIX(wt)
V5-pIX(1-20_S/T>A)
V5-pIX(21-40_S/T>A)
V5-pIX(41-60_S/T>A)
V5-pIX(61-80_S/T>A)
V5-pIX(81-100_S/T>A)
V5-pIX(101-140_S/T>A)
HA-Ubi - + + + + + + + +
Input
WB: HA
15-
WB: V5
IP
70-
55-
35-
25-
4xUbi
3xUbi
2xUbi
1xUbi
WB: HA
15-
V5-pIX
WB: V5
D
Input
IP (V5)
V5-pVII + +
V5-pIX + +
HA-Ubi + + + + + +
55-
35-
25-
4xUbi
3xUbi
2xUbi
1xUbi
WB:HA
25-
15-
V5-pVII
V5-pIX
WB:V5
E
Input
IP (HA)
pXI(2Y>A)-HA + +
pIX(wt)-HA + +
V5-Ubi + + + + + +
55-
35-
25-
4xUbi
3xUbi
2xUbi
1xUbi
WB:V5
15-
*
pIX-HA
WB:HA

### Fig. S3

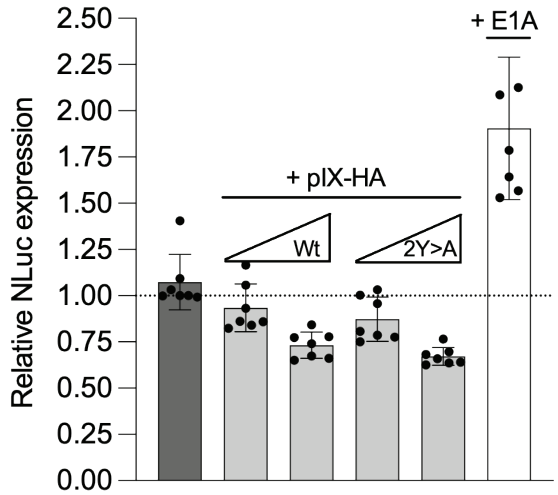

Relative NLuc expression
2.50
2.25
2.00
1.75
1.50
1.25
1.00
0.75
0.50
0.25
0.00
+ pIX-HA
Wt
2Y>A
+ E1A


**Fig. S4**

#### Supplementary material and methods

##### Plasmid constructions

Synthetic HAdV-C5 pIX and 19K/IX cDNA fragments and their respective mutants (Twist Biosciences) were cloned into a pcDNA3 vector in-frame with the N-terminal Flag (Flag-pIX, Flag-19K/IX), HA (HA-pIX), V5 (V5-pIX, V5-19K/IX), or C-terminal HA (pIX-HA) antibody epitope tags (1, 2). Note that we used 19K/IX cDNA, not the E1B genomic region, to generate the different epitope-tagged 19K/IX expression constructs. The plasmids expressing PSMC3-Flag and PSMD9-Flag were constructed by cloning synthetic genes (NM_002804.4 and NM_002813.7, Twist Biosciences) into the pcDNA(C-Flag) vector background. Plasmid pRK5-HA-Ubiquitin was purchased from Addgene (#17608); the His-Ubiquitin-expressing plasmid has been described before (3). HA-Ubiquitin mutants K11R and K48R were generated by cloning the respective mutation-containing DNA fragment into the pRK5-HA-Ubiquitin vector background. The NanoLuc (pNL-pIX_promoter) reporter plasmid was generated by cloning the pIX promoter (4), corresponding to nucleotides 3510-3613 of HAdV-C5 (AC_000008.1), into the pNL1.1.PGK[*Nluc*/PGK] vector (Promega) using the KpnI and NcoI restriction enzymes. The HAdV-C5 E1A transcription unit (nucleotides 1-1740; AC_000008.1) was cloned into the pUC19 plasmid using the SalI and EcoRI restriction enzymes to express E1A proteins (pUC-E1A).

##### Isolation of the Adenobuilder viruses

Five days after transfection with assembled Adenobuilder genomes, cells underwent three freeze/thaw cycles to release viral particles. The resulting lysate was then used to infect HEK293AD or H1299 cells to amplify the virus. After 3 to 5 days of infection, the cells were collected, resuspended in 400 µl of cell growth medium, and lysed through three freeze/thaw cycles on dry ice and at +37°C. Cell lysates were centrifuged at 5000 × g for 5 min at +4°C, and the virus-containing supernatants were aliquoted and stored at -80°C.

**Virus titration**

A549 cells were seeded into a 24-well plate and infected with virus dilutions ranging from $10^{-2}$ to $10^{-6}$. At 40 hpi, cells were fixed in 4% paraformaldehyde (PFA) in Phosphate-Buffered Saline (PBS), then permeabilized with 0.125% Triton X-100/PBS (1). The cells were blocked in 2% BSA/PBS for 1 h at room temperature, then incubated overnight at +4°C with an anti-hexon antibody (TC31-9C12.C9-s, Developmental Studies Hybridoma Bank (DSHB), Table S3). Infected cells were detected with a fluorescently labeled secondary antibody (Alexa Fluor 488, A11001; Thermo Fisher Scientific). Fluorescent forming units (FFUs) were manually counted from appropriately diluted samples using a Nikon Eclipse TS100 microscope equipped for fluorescence detection at 10× magnification. Final virus titers (FFU/ml) were calculated using the dilution factor and the number of positively stained cells per well in a 24-well plate.

**cDNA synthesis and qRT-PCR**

Complementary DNA (cDNA) was generated using the Maxima reverse transcriptase (Thermo Fisher Scientific) primed with random hexamer primers (Thermo Fisher Scientific, SO142). Quantitative RT-PCR (qRT-PCR) reactions were prepared with FIREPol EvaGreen qPCR Supermix (Solis Biodyne, 08-36-00020) and amplified in a QuantStudio 6 Flex real-time PCR system (Thermo Fisher Scientific). Data from a triplicate experiment were normalized to housekeeping genes (HRPT1), and relative mRNA expression (mean ± SD) was calculated using the ΔΔCt method. Primer sequences are available in Table S4.

**Mass spectrometry**

Analysis of MS raw files was performed using the FragPipe quantitative proteomics software suite (version 24.0), employing MSFragger (version 4.4.1) (5) for database searching and IonQuant (version 1.11.11) (6) for label-free quantification (LFQ). The LFQ-MBR (match-between-runs) workflow was used. Spectra were searched against the UniProtKB/Swiss-Prot *Homo sapiens* proteome (UP000005640, version 2024-11-21) and UniProtKB/Swiss-Prot human adenovirus C serotype 5 (UP000004992, version 2026-03-27) databases together with an in-house contaminant database. The precursor mass tolerance was set to 20 ppm, and the fragment mass tolerance to 0.6 Da. Up to two missed tryptic cleavages were allowed. Methionine oxidation and protein N-terminal acetylation were specified as variable modifications, and cysteine carbamidomethylation as a fixed modification. Peptide-spectrum matches (PSM) generated by MSFragger were rescored with MSBooster (version 1.4.14) (7) using AlphaPeptDeep-based prediction models for fragment ion spectra (AlphaPept MS2 Generic) and retention time (AlphaPept RT Generic) (8), followed by statistical validation with Percolator (version 3.7.1) (9). Protein inference was performed with ProteinProphet, and false discovery rate (FDR) filtering and report generation were carried out within the Philosopher-based FragPipe workflow (version 5.1.3-RC9) (10), with identifications filtered to 1% at the PSM, peptide, and protein levels. Downstream analysis was performed in FragPipe-Analyst (version 1.23) (11) using $\log_2$-transformed IonQuant protein intensities after removal of potential contaminant proteins. Differential enrichment (DE) analysis was carried out using the limma framework, which fits feature-wise linear models and applies empirical Bayes moderation of the standard errors to obtain moderated *t*-statistics and associated *P*-values. Variance-stabilizing normalization (vsn) was applied. Missing values were imputed using Perseus-type imputation, in which missing values are replaced by random values drawn from a normal distribution with a downshift of 1.8 standard deviations and with a width of 0.3 for each sample. *P*-values were adjusted for multiple testing using the Benjamini-Hochberg method, and proteins with a $\log_2$ fold change of ≥2, and an adjusted *P*-value ≤0.05 were considered significantly enriched.

**NanoLuc reporter assay**

A549 cells grown in a 96-well plate were co-transfected with the pNL-pIX_promoter plasmid and increasing concentrations (20 ng and 80 ng) of the pIX(wt)-HA or pIX(Y2>A)-HA expressing plasmids. The total plasmid DNA concentration was kept the same (100 ng) in all the reactions with the empty pcDNA plasmid. E1A-expressing plasmid (pUC-E1A, 20 ng per well) was used as a positive control for pIX promoter activity. At 24 h post-transfection, the cells were lysed using the NanoGlo Luciferase Assay System (Promega), and the NanoLuc signal was detected with the Spark 10M plate reader (Tecan).

#### Table S1. Cloned plasmids used in the study

| Name | Identifier | Purpose | Insert | Epitope tag |
|---|---|---|---|---|
| pcDNA-V5-19K/IX(Wt) | pTP907 | WB/IF/IP | 19K/IX(Wt) | N-terminal V5 |
| pcDNA-Flag-pIX(Wt) | pTP886 | WB/IF/IP | pIX(Wt) | N-terminal Flag |
| pcDNA-HA-pIX(Wt) | pTP902 | WB/IF | pIX(Wt) | N-terminal HA |
| pcDNA-V5-pIX(Wt) | pTP970 | WB/IF/IP | pIX(Wt) | N-terminal V5 |
| pcDNA-HA-pIX(2K>R) | pTP955 | WB | pIX(K99R/K132R) | N-terminal HA |
| pcDNA-V5-pIX(1-20_S/T>A) | pTP973 | WB/IP | pIX(1-20_S/T>A) | N-terminal V5 |
| pcDNA-V5-pIX(21-40_S/T>A) | pTP974 | WB/IP | pIX(21-40_S/T>A) | N-terminal V5 |
| pcDNA-V5-pIX(41-60_S/T>A) | pTP975 | WB/IP | pIX(41-60_S/T>A) | N-terminal V5 |
| pcDNA-V5-pIX(61-80_S/T>A) | pTP976 | WB/IP | pIX(61-80_S/T>A) | N-terminal V5 |
| pcDNA-V5-pIX(81-100_S/T>A) | pTP977 | WB/IP | pIX(81-100_S/T>A) | N-terminal V5 |
| pcDNA-V5-pIX(101-140_S/T>A) | pTP978 | WB/IP | pIX(101-140_S/T>A) | N-terminal V5 |
| pcDNA-V5-pIX(2Y>A) | pTP980 | WB/IP | pIX(Y14A/Y49A) | N-terminal V5 |
| pcDNA-V5-pIX(Y49A) | pTP1003 | WB/IP | pIX(Y49A) | N-terminal V5 |
| pcDNA-V5-19K/IX(2Y>A) | pTP1002 | WB/IP | 19K/IX(Y77A/Y112A)* | N-terminal V5 |
| pcDNA-pIX(Wt)-HA | pTP1018 | WB/IF/IP/NLuc | pIX(Wt) | C-terminal HA |
| pcDNA-pIX(2Y>A)-HA | pTP1022 | WB/IF/IP/NLuc | pIX(Y14A/Y49A) | C-terminal HA |
| pcDNA-pIX(Y14A)-HA | pTP1029 | IF | pIX(Y14A) | C-terminal HA |
| pcDNA-pIX(Y49A)-HA | pTP1030 | IF | pIX(Y49A) | C-terminal HA |
| pcDNA-PSMC3-Flag | pTP1015 | WB/IF | PSMC3 | C-terminal Flag |
| pcDNA-PSMD9-Flag | pTP1016 | WB/IF | PSMD9 | C-terminal Flag |
| pRK5-HA-Ubi(Wt) | pTP994 | WB/IP | Ubiquitin(Wt) | N-terminal HA |
| pRK5-HA-Ubi(K11R) | pTP1043 | WB/IP | Ubiquitin(K11R) | N-terminal HA |
| pRK5-HA-Ubi(K48R) | pTP1044 | WB/IP | Ubiquitin(K48R) | N-terminal HA |
| pRK5-HA-Ubi(K11R/K48R) | pTP1045 | WB/IP | Ubiquitin(K11R/K48R) | N-terminal HA |
| pcDNA-V5-pVII(K26R/K27R) | pTP979 | WB/IP | pVII(K26R/K27R) | N-terminal V5 |
| pUC-E1A | pTP699 | NLuc | E1A(nt. 1-1740) | N/A |
| pNL-pIX_promoter | pTP1028 | NLuc | pIX (nt. 3510-3613) | N/A |

*The point mutations Y14A/Y49A in pIX correspond to amino acids Y77A/Y112A in the 19K/IX protein, due to 63-amino-acid N-terminal extension from E1B19K CDS.

WB; Western blot, IF; immunofluorescence, IP; immunoprecipitation, NLuc; NanoLuc reporter gene assay

#### Table S2. Adenobuilder-based viruses used in the study

| Virus name | Epitope tagged and modified virus genes |
|---|---|
| Wt_AB | unmodified AdenoBuilder-based (1) wild-type HAdV-C5 expressing no epitope-tagged proteins |
| Wt | **Flag**-E1B19K<br>pIX-**HA**<br>**Flag**-19K/IX-**HA** |
| Stop$^M$ | **Flag**-E1B19K<br>pIX-**HA** |
| 5'ss$^M$ | **Flag**-E1B19K<br>pIX-**HA** |
| Wt_Gal4 | **Flag**-E1B19K<br>pIX-**HA**<br>**Flag**-19K/IX-**HA**<br>**V5**-Gal4 |
| Stop$^M$_Gal4 | **Flag**-E1B19K<br>pIX-**HA**<br>**V5**-Gal4 |
| Stop$^M$_V5-19K/IX | **Flag**-E1B19K<br>pIX-**HA**<br>**V5**-19K/IX |
| Wt(2K>R) | **Flag**-E1B19K<br>pIX(K99R/K132R)-**HA**<br>**Flag**-19K/IX(K162R/K195R)-**HA*** |
| Stop$^M$(2K>R) | **Flag**-E1B19K<br>pIX(K99R/K132R)-**HA** |
| Wt(2Y>A) | **Flag**-E1B19K<br>pIX(Y14A/Y49A)-**HA**<br>**Flag**-19K/IX(Y77A/Y112A)-**HA*** |
| Stop$^M$ (2Y>A) | **Flag**-E1B19K<br>pIX(Y14A/Y49A)-**HA** |

***** The point mutations in pIX are also present in the 19K/IX protein, which contains a 63-amino-acid N-terminal extension.

1. Jang Y, Bunz F. 2022. AdenoBuilder: A platform for the modular assembly of recombinant adenoviruses. Star Protocols 3.

Table S3. Antibodies used for western blot, immunofluorescence, and virus titer assays

| Target | Dilution | Purpose | Host | Source | Identifier/reference |
|---|---|---|---|---|---|
| Anti-pV | 1:5000 | WB | Rabbit | D.Matthews | (1) |
| Anti-E1B55K (2A6) | 1:40 | WB | Mouse | T.Dobner | (2) |
| Anti-AVP | 1:100 | WB | Rabbit | M.Balakirev | (3) |
| Anti-DBP | 1:500 | IF | Mouse | A.Levine | A72K A1-6, (4) |
| Anti-GAPDH (FL-335) | 1:1000 | WB | Rabbit | Santa Cruz | sc-25778 |
| Anti-Tubulin (6A204) | 1:1000 | WB | Mouse | Santa Cruz | sc-69969 |
| Anti-Gal4 DBD | 1:1000 | WB | Mouse | Santa Cruz | sc-510 |
| Anti-E1A(M58) | 1:1000 | WB/IF | Mouse | Santa Cruz | sc-58658 |
| Anti-V5 | 1:1000 | WB/IF | Rabbit | Proteintech | 14440-1-AP |
| Anti-CoxIV | 1:1500 | WB | Rabbit | Proteintech | 11242-1-AP |
| Anti-HA | 1:1000 | WB/IF | Rabbit | Proteintech | 51064-2-AP |
| Anti-Flag | 1:1000 | WB/IF | Rabbit | Proteintech | 12552-1-AP |
| Anti-PSMC3 | 1:1000 | WB | Rabbit | Proteintech | 24142-1-AP |
| Anti-PSMD9 | 1:1000 | WB | Rabbit | Proteintech | 26922-1-AP |
| Anti-Flag | 1:1000 | WB/IF | Mouse | Proteintech | 66008-4-IG |
| Anti-V5 | 1:1000 | WB | Mouse | Proteintech/Chromotek | SV5-P-K |
| Anti-Adenovirus type 5 | 1:2000 | WB | Rabbit | Abcam | Ab6982 |
| Anti-Histone H3 | 1:30.000 | WB | Rabbit | Abcam | ab1791 |
| Anti-HSPA1A | 1:1000 | WB | Mouse | Thermo | MA3-006 |
| Anti-His | 1:1000 | WB | Mouse | Takara | 631212 |
| Anti-Hexon | 1:100 | Titration | Mouse | DSHB | TC21-9C12-C9 |
| Anti-Fiber | 1:2000 | WB | Mouse | Invitrogen | MA5-11222 |
| Anti-HA.11 | 1:1000 | WB/IF | Mouse | BioLegend | 901513 |
| Anti-HA | 1:1000 | WB | Rabbit | Sigma | H6908 |

WB; Western blot, IF; immunofluorescence, Titration; virus titration

#### Table S4. PCR/qPCR primers to detect host cell and viral mRNAs

| **Target** | **Forward** | **Reverse** | **Purpose** |
|---|---|---|---|
| HAdV-C5 Flag-19K/IX-HA | 5'-GCG*GAATTC*GCCACC**ATGGACTACAAGG**-3' | 5'-TGC*CTCGAG*TCATTA**AGCGTAATC**-3' | Viral mRNA detection by endpoint PCR |
| GAPDH | 5'-GGTTTACATGTTCCAATATGATTCCA-3' | 5'-ATGGGATTTCCATTGATGACAAG-3' | Housekeeping gene mRNA detection by endpoint PCR |
| HRPT1 | 5'- TGGACAGGACTGAACGTCTTG -3' | 5'- CCAGCAGGTCAGCAAAGAATTTA-3' | Housekeeping gene mRNA detection by qRT-PCR |
| HAdV-C5 E1A | 5'- GTGCCCCATTAAACCAGTTG-3' | 5'- GGCGTTTACAGCTCAAGTCC-3' | Viral mRNA detection by qRT-PCR |
| HAdV-C5 pIX | 5'-CCGCGATGACAAGTTGACGG-3' | 5'-CATTGGGAGGGGAGGAAGCC-3' | Viral mRNA detection by qRT-PCR |

Primer 5’-GCG*GAATTC*GCCACC**ATGGACTACAAGG**-3' contains an EcoRI restriction site (Italics) and a sequence complementary to the Flag antibody epitope-coding sequence (bold)

Primer 5'-TGC*CTCGAG*TCATTA**AGCGTAATC**-3' contains an XhoI restriction site (Italics) and a sequence complementary to the HA antibody epitope-coding sequence (bold)

The primer pair (5′-CCGCGATGACAAGTTGACGG-3′ and 5′-CATTGGGAGGGGAGGAAGCC-3′) amplifies pIX CDS corresponding to nucleotides 3893–4023 of the HAdV-C5 genome.

The primer pair (5’-GCG*GAATTC*GCCACC**ATGGACTACAAGG**-3' and 5'-TGC*CTCGAG*TCATTA**AGCGTAATC**-3') amplifies only the Flag-19K/IX-HA transcript and not the Flag-E1B19K or pIX-HA transcripts.
